# Starvation-independent alarmone production inhibits translation through GTP depletion

**DOI:** 10.64898/2026.01.12.699007

**Authors:** Kevin A. England, Aude Trinquier, Leah K. McKinney, Jue D. Wang, Heather A. Feaga

## Abstract

Bacteria produce the alarmone nucleotides (p)ppGpp during stress to affect replication, transcription, translation, and metabolism. Recently, pGpp was identified as a third alarmone that is produced from the hydrolysis of (p)ppGpp. Although pGpp is a major component of bacterial stress responses, its precise role in mediating these responses is poorly understood. ppGpp and pppGpp bind translation GTPases and therefore directly affect translation to conserve resources during periods of stress. Here, we show that while pGpp is a weaker inhibitor of protein synthesis than ppGpp and pppGpp *in vitro*, pGpp production in the model Gram-positive bacterium *Bacillus subtilis* leads to faster translation inhibition *in vivo*. Faster translation inhibition is accompanied by greater levels of disengaged ribosomal subunits and hibernating ribosome dimers, suggesting that translation initiation is strongly inhibited. We show that alarmone production *in vivo* causes a severe depletion of GTP, which is sufficient for translation inhibition. Finally, we find that pGpp production also causes more robust transcriptome remodeling than (p)ppGpp production. This work supports a model that implicates all three alarmones in translation inhibition *via* GTP depletion.

## Introduction

During stress, bacteria hyperphosphorylate guanosine nucleotides to produce the alarmones guanosine tetraphosphate (ppGpp) and guanosine pentaphosphate (pppGpp), collectively referred to as (p)ppGpp. These nucleotides invoke changes to replication, transcription, translation, metabolism, and other physiological pathways to mount an effective stress response (1–4). Through these mechanisms, (p)ppGpp production increases long-term survival and facilitates antibiotic persistence (5–7). Therefore, determining the molecular mechanisms by which (p)ppGpp production causes physiological changes is important for understanding how bacteria respond to their environment.

(p)ppGpp mounts a global transcriptional response to stress through various mechanisms. In Proteobacteria, (p)ppGpp binds RNA polymerase (RNAP) directly or *via* the transcription factor DksA to alter transcription elongation and initiation kinetics (8–10). Consequently, binding of (p)ppGpp to RNAP accounts for nearly all transcriptomic changes during alarmone production in *Escherichia coli* (11). However, (p)ppGpp does not bind to RNAP directly in most bacteria, including the model Gram-positive bacterium *Bacillus subtilis* (10, 12). Thus, an alternative mode for (p)ppGpp to drive a global transcriptomic response to stress is through GTP depletion. GTP depletion results from direct inhibition of many purine biosynthesis and salvage enzymes by alarmones (13–20), co-repression of the PurR regulon (19), and through concomitant consumption of GTP, GDP, or GMP by the active alarmone synthetases. Globally, GTP depletion reduces transcription of guanosine-initiating transcripts (including many exponential phase transcripts and all ten ribosomal RNA operons in *B. subtilis*) (12, 21) and reduces repression by CodY (a transcriptional repressor in the presence of GTP) (22–24).

(p)ppGpp has additional targets in the cell, and these include translational GTPases that act at initiation (IF2), elongation (EF-Tu and EF-G), ribosome recycling (EF-G and HflX), and ribosome maturation (e.g., Era, RsgA, RbgA) (13, 17, 18, 25–33). To date, there have been a few studies that characterize (p)ppGpp-mediated changes to translation *in vivo*. For example, alarmone synthesis in *B. subtilis* activates transcription of the gene encoding the ribosome hibernation promoting factor (HPF) (34), which dimerizes ribosomes that have disengaged from translation (35). Similarly, heat shock causes *B. subtilis* to produce (p)ppGpp, and this transient accumulation of (p)ppGpp is required to inhibit translation, induce expression of HPF and ribosome dimerization, and maintain 16S rRNA (36). Also in *B. subtilis*, during the transition to stationary phase, (p)ppGpp causes heterogeneous translation inhibition through interaction with the initiation factor IF2 (33); this is consistent with biochemical characterization showing that ppGpp exhibits strong binding affinity for IF2 (32).

Guanosine-5′-monophosphate-3′-diphosphate (pGpp) has recently been appreciated as a third alarmone. pGpp results from direct pyrophosphorylation of GMP by alarmone synthetases or by hydrolysis of (p)ppGpp (15, 17, 26, 35, 37–41). Upon alarmone induction in *B.* subtilis, the major source of pGpp is <u>N</u>uDiX <u>a</u>larmone <u>h</u>ydrolase <u>A</u> (NahA), which hydrolyzes (p)ppGpp to pGpp (26, 39). In contrast to (p)ppGpp, pGpp does not bind to translational GTPases with high affinity, neither in a differential radial capillary action of ligand assay (DRaCALA) screen using cell lysates from an open reading frame overexpression library (26) nor using pGpp capture compounds (25). For example, whereas ppGpp and pppGpp bind the translation elongation factor EF-G with micromolar affinity, no binding was detected between pGpp and EF-G, suggesting little to no affinity (26). Thus, it has been proposed that bacteria fine-tune translation inhibition during alarmone production by altering the concentrations of pGpp and (p)ppGpp. However, the isolated effects of both pGpp and (p)ppGpp on translation, independent of starvation or other stress, is largely unexplored.

Here, we determine the effects of alarmones on cell physiology using genetic induction of alarmone synthetases. This approach enables isolation of the effects of alarmone production alone by uncoupling alarmone production from exogenous starvation or stress. We genetically induce alarmone production in *B. subtilis* in the presence or absence of NahA and decipher differences between pGpp and (p)ppGpp on translation and global gene expression. We find that pGpp is a weaker inhibitor of protein synthesis than (p)ppGpp *in vitro*, consistent with its lesser affinity for translational GTPases (26). However, genetic induction of pGpp synthesis *in vivo* causes more rapid translation inhibition than (p)ppGpp synthesis. We then show that GTP depletion–which occurs concomitantly to alarmone production–is sufficient for translation inhibition. Lastly, we show that pGpp production leads to greater and more rapid changes in gene expression than (p)ppGpp production. This work highlights how production of different alarmone species can have differing effects on cell physiology and supports a model that implicates all three alarmones in translation inhibition *via* GTP depletion.

## Results

### (p)ppGpp inhibits protein synthesis more strongly than pGpp *in vitro*

(p)ppGpp binds translational GTPases that act at every step of translation, but pGpp binding to translational GTPases has not been detected (26). To test whether pGpp affects translation *in vitro*, we performed *in vitro* transcription-translation assays using the PURExpress system (NEB). In this reaction, we expressed DHFR under the control of a T7 promoter since T7 RNA polymerase is not inhibited by (p)ppGpp (33, 42). ATP and GTP starting concentrations are 2 mM, and ATP is replenished by adenylate kinase and creatine kinase and GTP is replenished by nucleoside diphosphate kinase; nevertheless, ATP is eventually exhausted, which quenches protein production after 1-2 hours (43). At the start of the reaction, we added increasing concentrations of purified pGpp, ppGpp, or pppGpp and measured DHFR production after incubating for 2 hours (Figure 1A).

**Figure 1.**
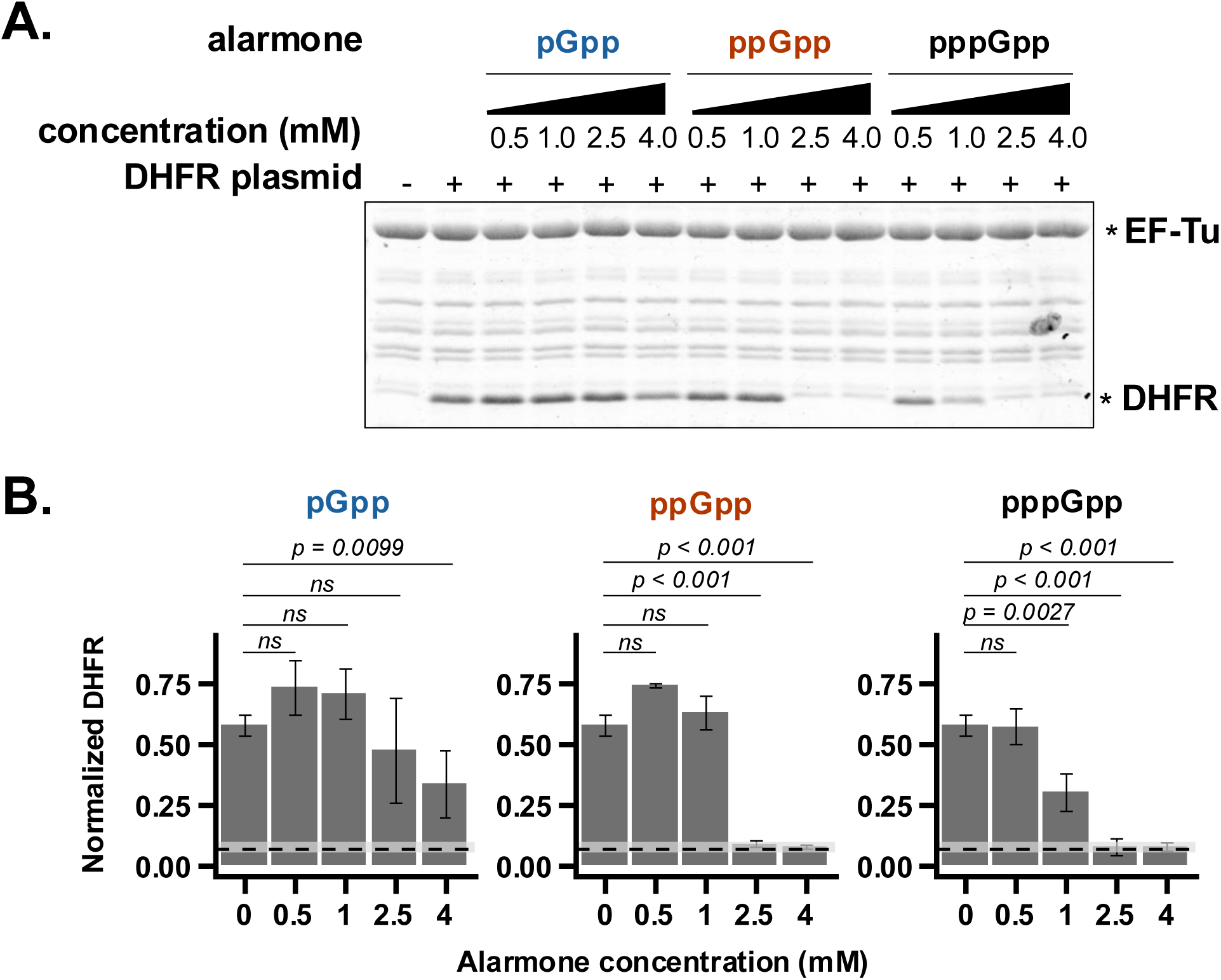
pGpp is a weaker inhibitor of protein synthesis *in vitro* than ppGpp and pppGpp. **(A)** Representative Coomassie-stained SDS-PAGE, showing the effect of increasing concentrations of purified pGpp, ppGpp, and pppGpp on production of DHFR (NEB PURExpress). **(B)** *In vitro* translation reactions densitometrically quantified from (A). Y-values represent mean DHFR normalized to EF-Tu as a loading control (+/- S.D., *n* = 3). Horizontal dashed lines and light gray boxes represents the mean EF-Tu-normalized background band intensity (+/- S.D.) from reactions lacking DHFR plasmid, which represents the limit of detection. A linear mixed-effects model was used, with fixed effects of alarmone, concentration, and their interaction and a random effect of replicate/batch. *P-*values represent results from a post-hoc Dunnett’s test to compare each concentration to the 0 mM control within each alarmone group.

The effect of alarmone concentration on DHFR production is significantly different for pGpp, ppGpp, and pppGpp (*F*_8,28_ = 5.97, *p* < 0.0002). Of the three alarmones, pGpp exhibits the least inhibition of protein synthesis (Figure 1B), with reduced DHFR at 4 mM pGpp, consistent with its lack of affinity for translational GTPases (25, 26). Meanwhile, ppGpp inhibited DHFR production to a much greater extent, with no detectable DHFR produced at 2.5 mM ppGpp, consistent with previous work that has determined ppGpp to be a potent inhibitor of this *in vitro* system (28, 33). Lastly, we found pppGpp to be the strongest inhibitor of DHFR production, with reduced DHFR at 1 mM pppGpp. These results indicate that pGpp is a weak inhibitor of *in vitro* protein synthesis, followed by ppGpp and pppGpp as stronger inhibitors.

### Starvation-independent production of pGpp and (p)ppGpp rapidly reduce GTP *in vivo* and arrest growth

Since pGpp is the weakest inhibitor of protein synthesis *in vitro*, we next determined the effects of pGpp and (p)ppGpp on translation *in vivo*. We used genetic induction of alarmone synthesis to isolate the effects of these alarmone species on translation because–unlike the use of amino acid starvation or other stressors–genetic induction does not directly change the concentrations of other translational substrates in the cell such as aminoacylated tRNAs. Thus, we created a pGpp inducible mutant and (p)ppGpp inducible mutant. In both mutants, all endogenous alarmone synthetases are deleted (both small synthetases, *sasA* and *sasB*, and the bifunctional (pp)pGpp synthetase/hydrolase *rel*) and a xylose-inducible *sasA* is integrated on the chromosome; the difference between the mutants is the presence or absence of the (p)ppGpp hydrolase *nahA*, respectively. Therefore, upon induction of SasA, the predominant guanosine alarmone is either pGpp in cells containing *nahA* or (p)ppGpp in Δ*nahA* cells as shown previously in similar genetic backgrounds (35, 39). In the absence of an inducer for SasA production, the pGpp and (p)ppGpp inducible strains grow comparable to wildtype; however, upon adding xylose during mid-exponential growth to induce *sasA* expression and alarmone production, growth is arrested within 40 minutes (Figure 2A).

**Figure 2.**
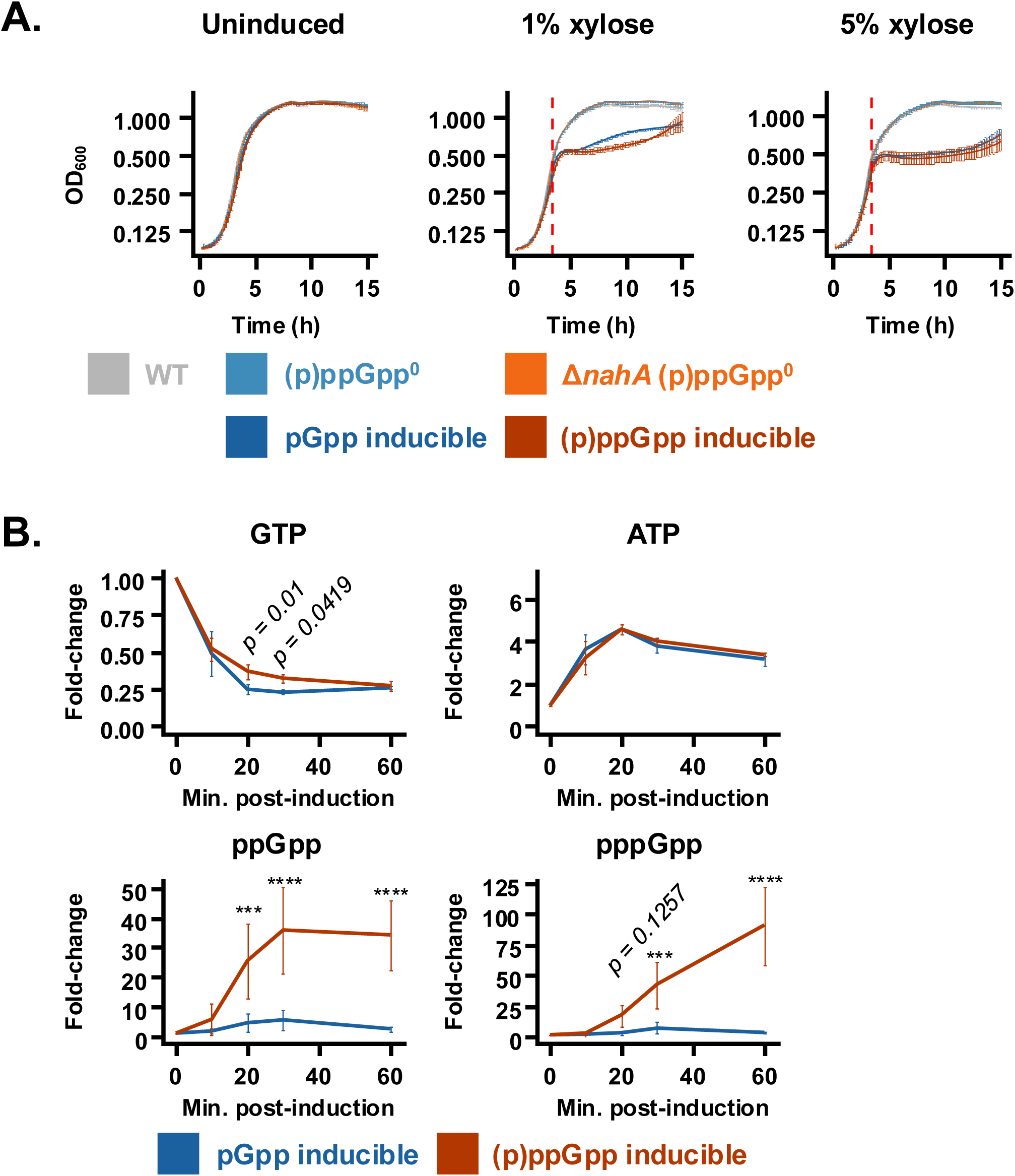
Starvation-independent production (i.e., genetic induction) of pGpp and (p)ppGpp *in vivo* arrests growth, reduces GTP, and increases ATP. **(A)** Mean growth curves (+/- S.D., *n* = 3) of pGpp and (p)ppGpp inducible strains, non-inducible parent strains, and wild-type in the absence of induction (left) or with 1% xylose (middle) or 5% xylose (right) added during exponential phase (red vertical dashed line). **(B)** Mean fold-change (vs. pre-induction) of GTP, ATP, ppGpp, and pppGpp after induction of pGpp and (p)ppGpp synthesis (+/- S.D., *n* = 3). Cultures were labeled with ^32^P-orthophosphate, and extracted nucleotides were resolved with thin-layer chromatography. A linear mixed-effects model was used, with time, strain, and their interaction as categorical fixed effects and replicate/batch a random effect. *P*-values represent results from pairwise comparisons between strains at each time-point.

Since pGpp and (p)ppGpp inhibit many enzymes involved in purine biosynthesis and salvage, we performed radiolabeling of nucleotides *in vivo* using ^32^P-orthophosphate and quantified changes in GTP and ATP using thin-layer chromatography (TLC). After just 10 minutes of inducing pGpp or (p)ppGpp synthesis, GTP decreases approximately 50%, and ATP increases approximately 3.4-fold (Figure 2B). GTP continues to decrease throughout alarmone induction, where it is depleted approximately 74% after 60 minutes, which is consistent with previous metabolomic measurements in similar genetic contexts (35, 39). Notably, at intermediate time-points, pGpp production depletes GTP modestly stronger than (p)ppGpp production (75% vs. 63% reduction at 20 minutes and 77% vs. 68% reduction at 30 minutes, respectively).

We also quantified changes in ppGpp and pppGpp after alarmone induction. In the (p)ppGpp inducible strain, ppGpp accumulates beyond basal levels by 20 minutes post-induction and reaches a maximum fold-change of 35.8 at 30 minutes post-induction (Figure 2B). pppGpp accumulates beyond basal levels slightly later than ppGpp, at 30 minutes post-induction, where it then increases to 90-fold at 60 minutes post-induction. In contrast, neither pppGpp nor ppGpp accumulate beyond basal levels in the pGpp inducible strain. Complementation of *nahA* to the (p)ppGpp inducible strain restores (p)ppGpp hydrolysis (Supplemental Figure S1). These results are consistent with previous metabolomics showing that NahA largely hydrolyzes (p)ppGpp into pGpp during SasA overexpression (39).

### pGpp production inhibits translation more rapidly than (p)ppGpp production *in vivo*

We next compared levels of translation inhibition between pGpp induction and (p)ppGpp induction *in vivo*. We incubated cells with the methionine analog *L*-homopropargylglycine (HPG) to measure total protein synthesis pre- and post-alarmone induction (44). Total protein synthesis is rapidly reduced upon (p)ppGpp induction and continues to decrease throughout induction (Figure 3A). Unexpectedly, pGpp induction also reduces total protein synthesis and does so more rapidly than (p)ppGpp induction. These results indicate that, despite being a weaker inhibitor of protein synthesis *in vitro*, *in vivo* pGpp production inhibits total protein synthesis more rapidly than (p)ppGpp production (Figure 3A).

**Figure 3.**
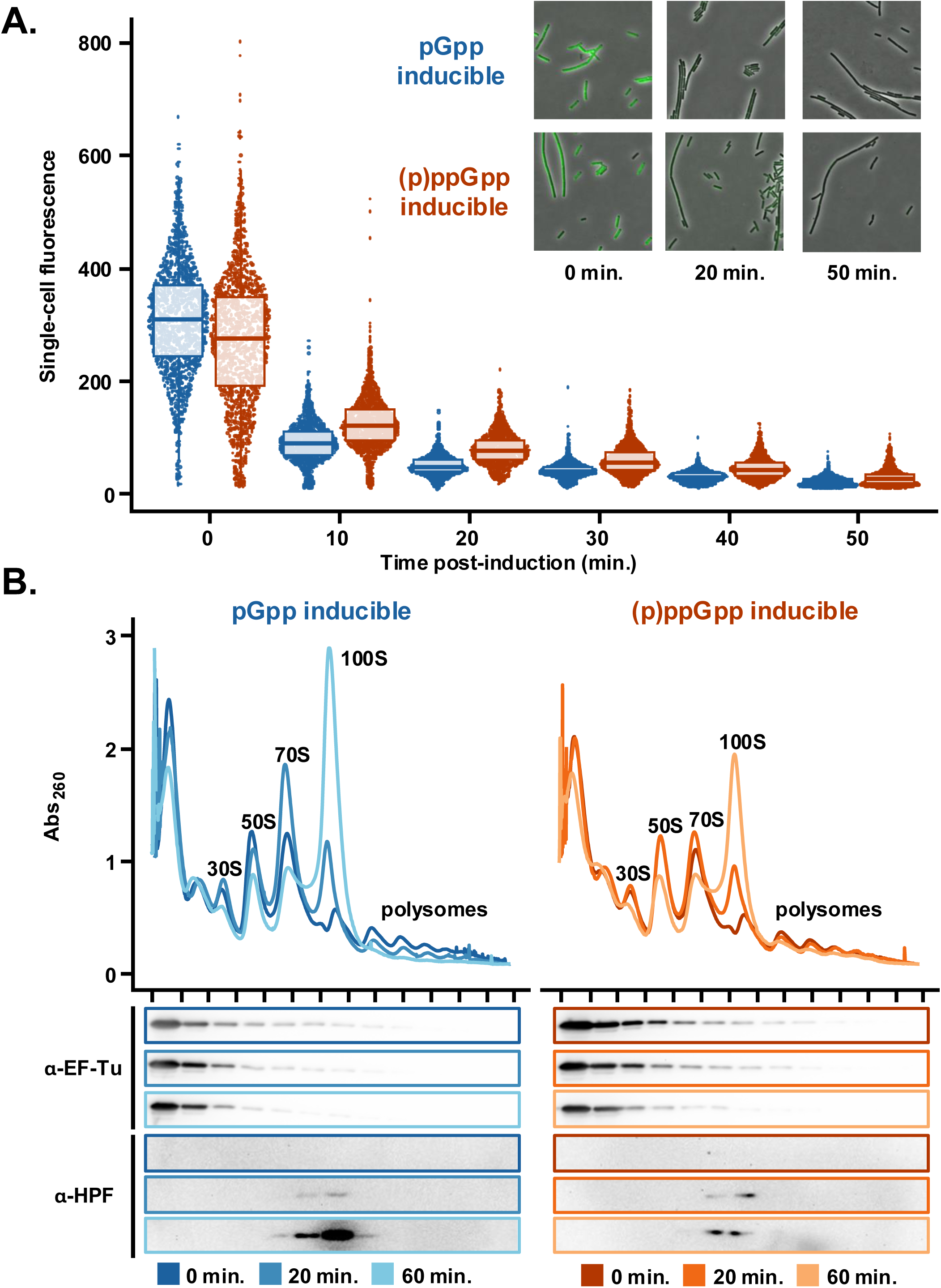
pGpp production inhibits translation faster than (p)ppGpp production. **(A)** Representative BONCAT to assess total protein synthesis pre- and post-pGpp and (p)ppGpp induction. **(B)** Representative polysome profiles of pGpp inducible and (p)ppGpp inducible mutants pre-, 20 minutes post-, and 60 minutes post-induction of alarmone, with gradient fraction western blotted with antibodies raised against EF-Tu or HPF.

As an additional method to monitor changes in protein synthesis, we examined changes in ribosome conformations during alarmone production using polysome profiling. Prior to alarmone induction, there are no detectable differences between the pGpp and (p)ppGpp inducible mutants (Figure 3B), suggesting that NahA does not greatly affect steady-state translation. Upon induction of alarmone synthesis, polysomes rapidly decrease. Polysomes are multiple ribosomes translating the same mRNA and, therefore, include highly translated transcripts (45). As a further measure of active translation, we probed gradient fractions with antibodies raised against the elongation factor EF-Tu. We observed reduced migration of EF-Tu into the gradient upon alarmone induction, consistent with fewer polysomes and translation inhibition. Concomitant to translation inhibition and a reduction in polysomes, HPF-bound 100S hibernating ribosome dimers increase (Figure 3B). Strikingly, pGpp production leads to a more rapid disappearance of heavy polysomes (i.e., 3 or more ribosomes translating the same mRNA) and a more rapid appearance of hibernating ribosome dimers than (p)ppGpp production (Supplemental Figure S2). Both the faster reduction in actively translating polysomes and the faster formation of hibernating ribosome dimers indicates that translation is more rapidly inhibited during pGpp production than (p)ppGpp production.

### GTP depletion is sufficient for translation inhibition and ribosome hibernation

While it was expected that (p)ppGpp production inhibits translation *in vivo* given its strong affinity for translational GTPases, it was unexpected that pGpp production also inhibits translation given its weak to no affinity for GTPases (Figure 3). Since we observed rapid GTP depletion during both pGpp and (p)ppGpp induction (Figure 2B), we hypothesized that genetic induction of alarmone synthesis inhibits translation through GTP depletion. To test whether GTP depletion alone is sufficient to inhibit translation, we used the GuaB/IMPDH inhibitor mycophenolic acid (MPA) (6) and identified a concentration that depletes GTP to a similar extent as genetic induction of pGpp or (p)ppGpp synthesis (Figure 4A). To avoid changes to basal pGpp and (p)ppGpp concentrations during MPA treatment that would confound the effects of GTP depletion alone, these experiments were performed in a (pp)pGpp^0^ background (Δ*sasB* Δ*sasA* Δ*rel*::*kan*). We then performed polysome profiling on MPA-treated cells to assess translation. MPA treatment causes a reduction in polysomes and a concomitant increase in 100S hibernating ribosome dimers (Figure 4B), consistent with what we observe during alarmone production. These results suggest that GTP depletion is sufficient for translation inhibition and ribosome hibernation.

**Figure 4.**
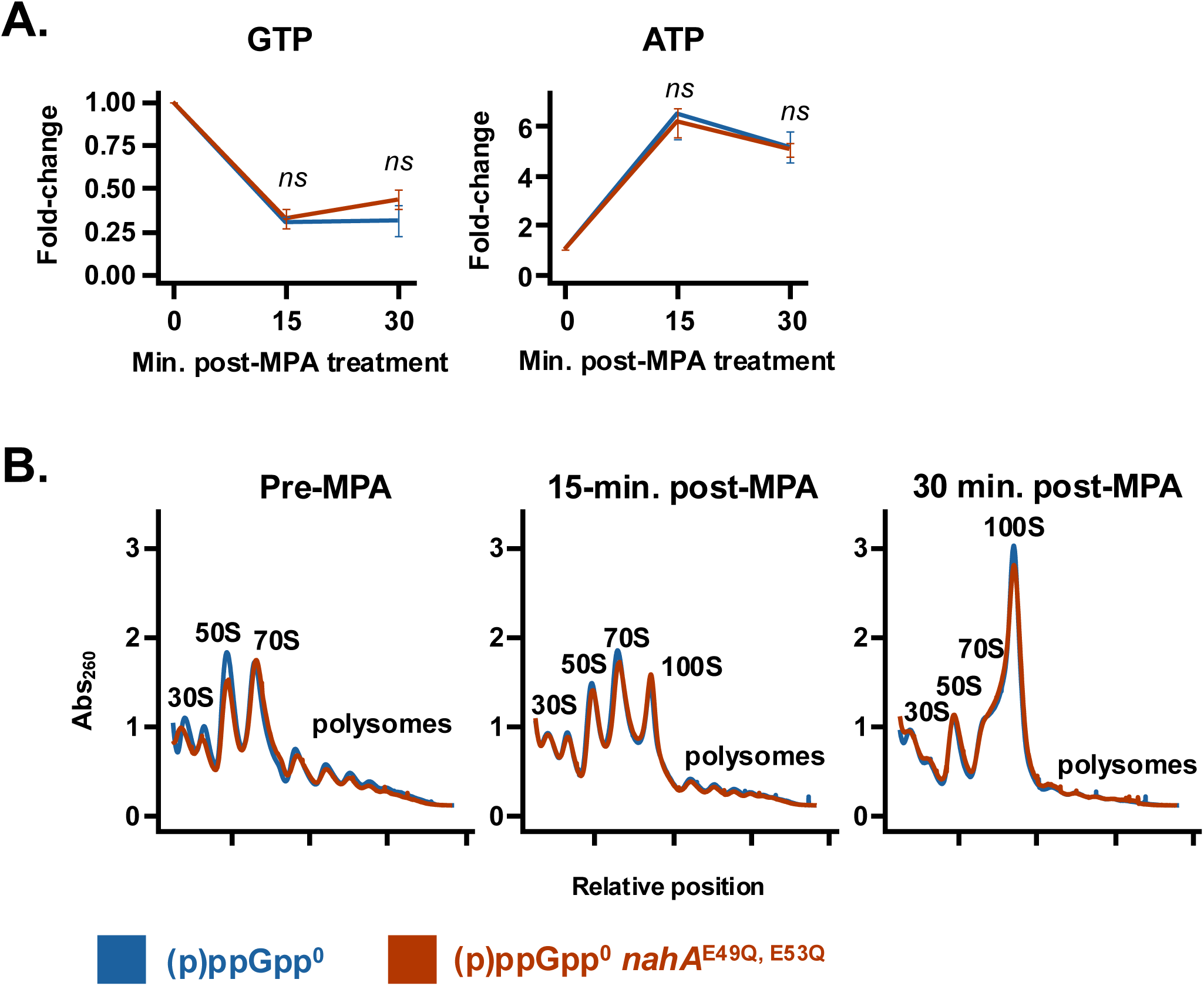
GTP depletion is sufficient for translation inhibition and ribosome dimerization, but not sufficient for the *nahA-*dependent enhancement. **(A)** Fold-change (vs. untreated) in GTP and ATP in (p)ppGpp^0^ and (p)ppGpp^0^ *nahA*^E49Q, E53Q^ mutants following mycophenolic acid (MPA) addition. Values represent means (+/-S.D., *n =* 3), and *p*-values are the result of an unpaired two-tailed Student’s T-test at each time-point. (**B)** Representative polysome profiles of (p)ppGpp^0^ and (p)ppGpp^0^ *nahA*^E49Q, E53Q^ mutants following MPA treatment.

To test whether NahA affects translation during GTP depletion in the absence of alarmone production, we depleted GTP from cells expressing a catalytically inactive NahA (NahA^E49Q, E53Q^) and lacking alarmones ((pp)pGpp^0^). Depleting GTP with MPA in (pp)pGpp^0^ *nahA*^E49Q, E53Q^ leads to a similar reduction in polysomes and increase in 100S hibernating ribosome dimers as cells expressing wildtype *nahA* (Figure 4A-B). These results suggest that NahA-mediated effects on translation are dependent on alarmone production.

### pGpp production causes ribosomes to disengage from translation more rapidly than (p)ppGpp production, independent of ribosome hibernation

In addition to a faster decrease in polysomes and a faster increase in hibernating ribosome dimers during pGpp production compared to (p)ppGpp production (Supplemental Figures S2), we also observed a transient increase in 70S monosomes only during pGpp production (Figure 3B). The lack of this transient accumulation of 70S monosomes during (p)ppGpp production can be restored by complementation with *nahA* to restore (p)ppGpp hydrolysis (Supplemental Figure S3). We hypothesized that these 70S ribosomes are disengaged from translation but not yet hibernating for two reasons. First, because there is no free HPF in the lightest gradient fractions (Figure 3B), HPF levels are likely non-saturating, which suggests that not all ribosomes disengaged from translation are dimerized. Second, at this time-point, heavy polysome levels are significantly lower during pGpp production compared to (p)ppGpp, yet the levels of hibernating ribosome dimers are equal (Supplemental Figure S2); these results suggest that ribosomes disengaged from translation are accumulating as both dimerized and non-dimerized species during pGpp production.

To quantify the total proportion of disengaged ribosomes and assess how engaged the 70S ribosomes are in translation, we performed polysome profiling in the presence of high concentrations of monovalent salt (KCl). High concentrations of monovalent salt specifically dissociate translationally disengaged ribosomes that lack a peptidyl-tRNA, are unbound to mRNA, or are hibernating dimers (46–53) into disengaged subunits. As a positive control to show that ribosomes lacking a peptidyl-tRNA are susceptible to dissociation by the high KCl concentrations used in our assay, we also included a set of samples where we added puromycin during cell lysis. Puromycin terminates translation and releases the nascent chain from the peptidyl-tRNA, which renders those ribosomes susceptible to dissociation by high-salt (51, 53).

Prior to alarmone induction, the pGpp and (p)ppGpp inducible strains exhibit similar levels of disengaged ribosomal subunits in polysome profiles, regardless of KCl concentration (Figure 5 left). As expected, the addition of KCl to sucrose gradients for polysome profiles increases the proportion of 30S and 50S ribosomal subunits and decreases the proportion 70S. Also, as expected, puromycin treatment increases the proportion of disengaged ribosomal subunits in polysome profiling in the presence of KCl. These results confirm that KCl dissociates vacant 70S ribosomes (representative polysome profiles are shown in Supplemental Figure S4A-C) and suggest that–in both strains prior to alarmone induction–55% of ribosomes are disengaged from translation.

**Figure 5.**
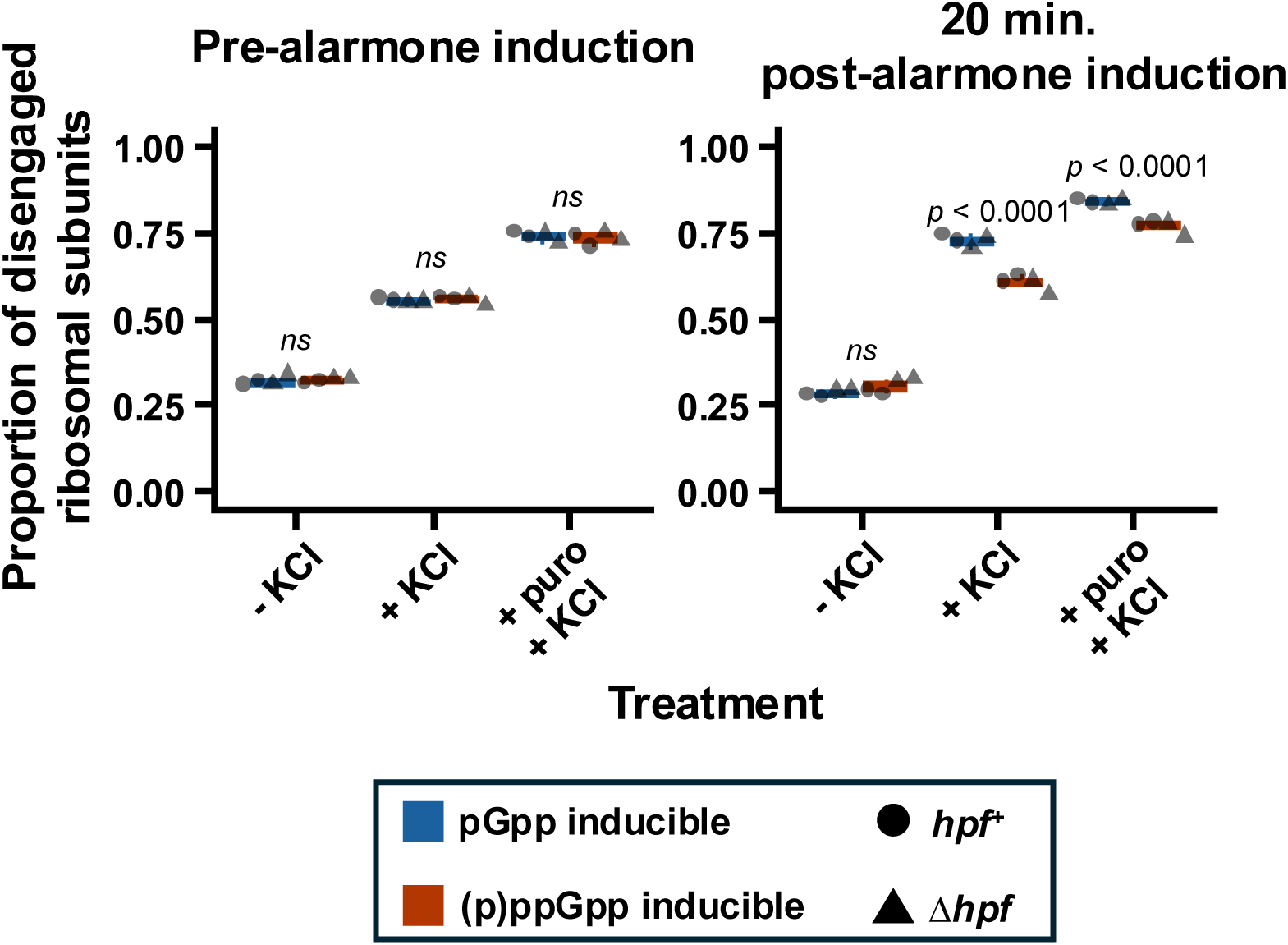
pGpp induction leads to faster disengagement of ribosomes, independent of dimer formation/hibernation. Cells were harvested for polysome profiling pre- and 20-minutes post-pGpp and (p)ppGpp synthesis, in the presence (*hpf*^+^) or absence (Δ*hpf*) of HPF. Polysome profiling was performed under normal salt conditions (no treatment), in the presence of 1 M KCl, or with puromycin treatment and 1 M KCl. KCl dissociates all disengaged ribosomes (lacking a peptidyl-tRNA and/or mRNA) into disengaged subunits, so that the total pool of disengaged ribosomes is equal to the proportion of subunits. A linear mixed-effects model with fixed effects of induction, alarmone, *hpf*, treatment, all two-way, three-way, and four-way interactions of those terms, and a random effect of replicate/batch was used. *P*-values represent results from pairwise comparisons of pGpp and (p)ppGpp within each level of induction and treatment, irrespective of *hpf*. Testing for the main effect of *hpf^+^* vs. Δ*hpf* across all levels of induction, treatment, and pGpp and (p)ppGpp yields *p* = 0.3611.

We next repeated the ribosome dissociation assays using lysates collected after 20 minutes of pGpp or (p)ppGpp induction. Under polysome profiling conditions lacking KCl, the proportion of disengaged ribosomal subunits are indistinguishable between the two strains and comparable to pre-induction conditions (Figure 5 right, Supplemental Figure S4D). However, polysome profiling with KCl reveals that 20 minutes of pGpp induction leads to significantly more disengaged ribosomes than (p)ppGpp induction (*p* < 0.0001). Importantly, polysome profiling in the presence of KCl also substantially dissociates 70S ribosomes (Supplemental Figure S4E); this indicates that the transient accumulation of 70S ribosomes post-induction of pGpp synthesis are mostly disengaged from translation as predicted. Incubating lysates with puromycin further increases the proportion of disengaged ribosomal subunits (Supplemental Figure S4F), albeit more for pGpp production than (p)ppGpp production (*p* = 0.0001). These results indicate that ribosomes become disengaged from translation more rapidly during pGpp production than (p)ppGpp production *in vivo*.

We previously observed that pGpp production leads to faster ribosome dimerization than (p)ppGpp production (Figure 3, Supplemental Figure S2). Therefore, we asked whether HPF-mediated dimer formation is sufficient to explain the more rapid disengagement of ribosomes upon pGpp induction. To test this, we performed the same ribosome dissociation assays in strains lacking HPF (Δ*hpf::tet*). Even in a Δ*hpf* background, ribosomes disengage from translation more rapidly during pGpp production than (p)ppGpp production (Figure 5 triangle symbols, Supplemental Figure S5). Additionally, ribosome dimerization does not affect the proportion of disengaged ribosomes (*p* = 0.3611, linear mixed-effects model). These results suggest that, regardless of ribosome dimerization, pGpp production causes ribosomes to disengage from translation more rapidly than (p)ppGpp production and that ribosome dimerization is a consequence, not a cause, of ribosomes becoming disengaged during alarmone induction.

### pGpp production remodels the transcriptome faster than (p)ppGpp production

One molecule of GTP is required for translation initiation by IF2, one molecule of GTP for every round of tRNA delivery by EF-Tu, and one molecule of GTP for every round of translocation by EF-G (54). Therefore, GTP depletion inhibits translation (as observed in Figure 4), at least in in part, by reducing the activity of translational GTPases. GTP depletion also causes transcriptome remodeling, which will further affect translation in the cell indirectly. To characterize transcriptional differences that could indirectly account for the translational differences between pGpp and (p)ppGpp production, we performed RNA-sequencing (RNA-seq) pre-alarmone induction and 10 and 20 minutes post-alarmone production (Supplemental Dataset S1).

First, hierarchical clustering appropriately clusters samples according to both strain and time post-induction on the X-axis, and genes with expectedly similar expression profiles (e.g., genes in the same operon) cluster together on the Y-axis (Supplemental Figure S6, Supplemental Dataset S2). These results indicate that the RNA-seq data is high-quality for assessing differential expression of genes across time within a single strain and across strains within a single time. Next, we grouped genes into various Clusters of Orthologous Genes (COGs) and asked which groups contain the most down or upregulated genes following pGpp or (p)ppGpp production. RNA-seq revealed global transcriptomic changes that are consistent with available literature (11, 35, 55). Translation, ribosomal structure and biogenesis and nucleotide metabolism and transport are the most down-regulated gene groups in response to alarmone production (Figure 6A). Amino acid metabolism and transport are among the most activated genes following alarmone production. Notably, pGpp production leads to more differentially expressed genes than (p)ppGpp production across most functional groups. Quantifying the total number of genes that are down- or up-regulated after pGpp and (p)ppGpp production reveals that pGpp production leads to more differentially expressed genes than (p)ppGpp production (Figure 6B).

**Figure 6.**
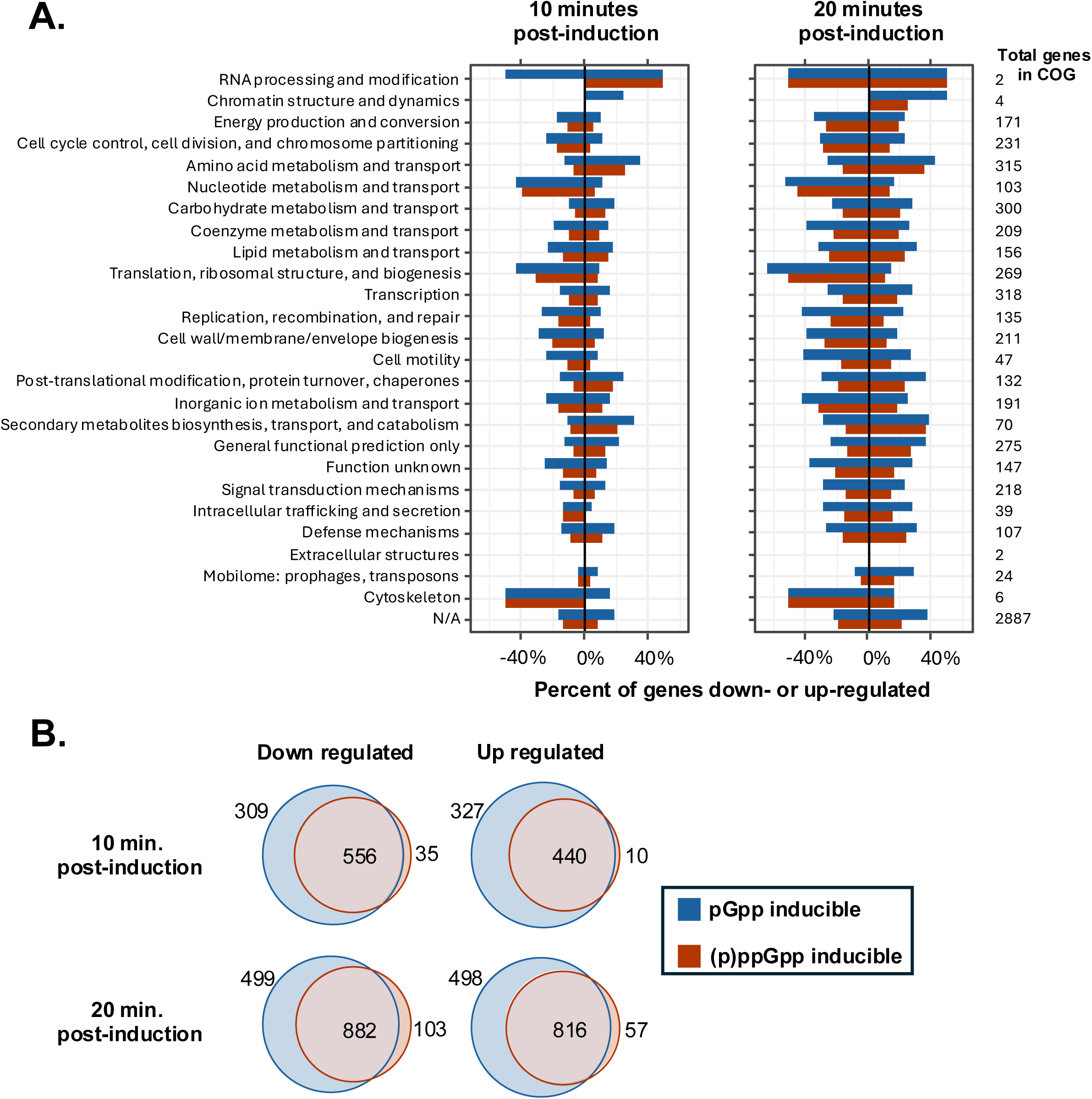
pGpp production remodels the transcriptome faster than (p)ppGpp production. **(A)** Percent of each Clusters of Orthologous Genes (COG) group that is differentially expressed post-induction of pGpp or (p)ppGpp synthesis (padj < 0.05 and |log_2_(fold change)| > 1), 10 minutes (left) and 20 minutes (right) post-induction. **(B)** Total number of genes that are down-or up-regulated, 10 minutes or 20 minutes post-induction pGpp or (p)ppGpp production.

Since pGpp production leads to a greater quantity of transcriptomic differences than (p)ppGpp production, we next examined whether pGpp production also leads to a greater degree of differential expression compared to (p)ppGpp production. To visualize this, we plotted the change in expression of all the genes that are differentially expressed during pGpp and/or (p)ppGpp production, where the X-coordinate is the change in expression during pGpp production and the Y-coordinate is the change in expression during (p)ppGpp production (Supplemental Figure S7). If changes in gene expression is equal during pGpp and (p)ppGpp production, then the points would fall along a line with an intercept of 0 and slope of 1. For example, comparing 20 minutes post-alarmone induction to pre-induction, *pyrG* encoding CTP synthase is significantly more repressed (log2FoldChange -1.40, *padj* = 2.77E-7) and *hpf* is more strongly activated (log2FoldChange 1.33, *padj* = 4.81E-16) by pGpp production than (p)ppGpp induction (Supplemental Dataset S1). In contrast, relative expression of *ilvB*, encoding the large subunit of acetolactate synthase and previously used as a reporter for low-GTP (6), increases equally during pGpp and (p)ppGpp induction. To assess whether pGpp production increases the degree of differential expression globally compared to (p)ppGpp induction, we transformed log2FoldChanges during pGpp or (p)ppGpp induction into the absolute value of log2FoldChanges and compared the means using paired *t*-test (*p* < 2.2E-16 at 10 minutes, and *p* < 2.2E-16 at 20 minutes post-induction). These results indicate that pGpp production evokes stronger changes to global gene expression than (p)ppGpp production.

### NahA does not affect the transcriptome without alarmone production

In addition to (p)ppGpp hydrolysis, NahA exhibits modest RNA pyrophosphohydrolysis *in vitro* (56). Pyrophosphohydrolase activity removes an ortho- or pyro-phosphate from the 5′ end of an mRNA, which primes it for recognition by the 5′-monophosphate-dependent exo- and endoribonucleases RNase J1 or RNase Y in *B. subtilis* (57–61). Therefore, we asked whether transcriptomic differences after alarmone production in the presence and absence of NahA (Figure 6) could be a consequence of differences in pyrophosphohydrolase activity, independent of the differences in pGpp and (p)ppGpp production. To investigate the possibility that NahA affects gene expression in the absence of alarmone production, we performed RNA-seq to compare the transcriptomes of cells expressing a wildtype NahA to cells expressing catalytically inactive NahA^E49Q, E53Q^ (56) in a (pp)pGpp^0^ background. Consistent with NahA having weak pyrophosphohydrolase activity as previously reported *in vitro* (56), NahA does not greatly affect the relative expression of any genes *in vivo* in the absence of alarmones (Supplemental Figure S8A).

To further explore whether NahA affects mRNA stability, we performed a rifampicin-sequencing (rif-seq) experiment to compare mRNA half-lives in (pp)pGpp^0^ mutants expressing wildtype or inactive NahA^E49Q, E53Q^. Rif-seq utilizes rifampicin to inhibit transcription initiation; performing RNA-seq before and after treatment then allows approximation of mRNA half-life. Consistent with NahA having poor pyrophosphohydrolase activity, NahA does not detectably destabilize mRNA globally (Supplemental Figure S8B, Supplemental Dataset S3). These data further support the conclusion that the transcriptomic differences between pGpp and (p)ppGpp production *in vivo* result as consequences of pGpp and (p)ppGpp production and not differences in mRNA pyrophosphohydrolase activity.

### The NuDiX motif of NahA is important for (p)ppGpp hydrolysis *in vitro*

Since the NuDiX motif of NahA is important for mRNA pyrophosphohydrolase activity *in vitro* (56) and we have determined that this activity does not strongly affect gene expression or mRNA stability *in vivo* (Supplemental Figure S8), we next tested whether this motif is important for (p)ppGpp hydrolysis. We performed *in vitro* hydrolysis of purified ppGpp and pppGpp using wildtype NahA and NahA^E49Q, E53Q^. We found that, while wildtype NahA readily hydrolyzes both alarmones as previously reported (26), NahA^E49Q, E53Q^ cannot hydrolyze ppGpp or pppGpp (Figure 7).

**Figure 7.**
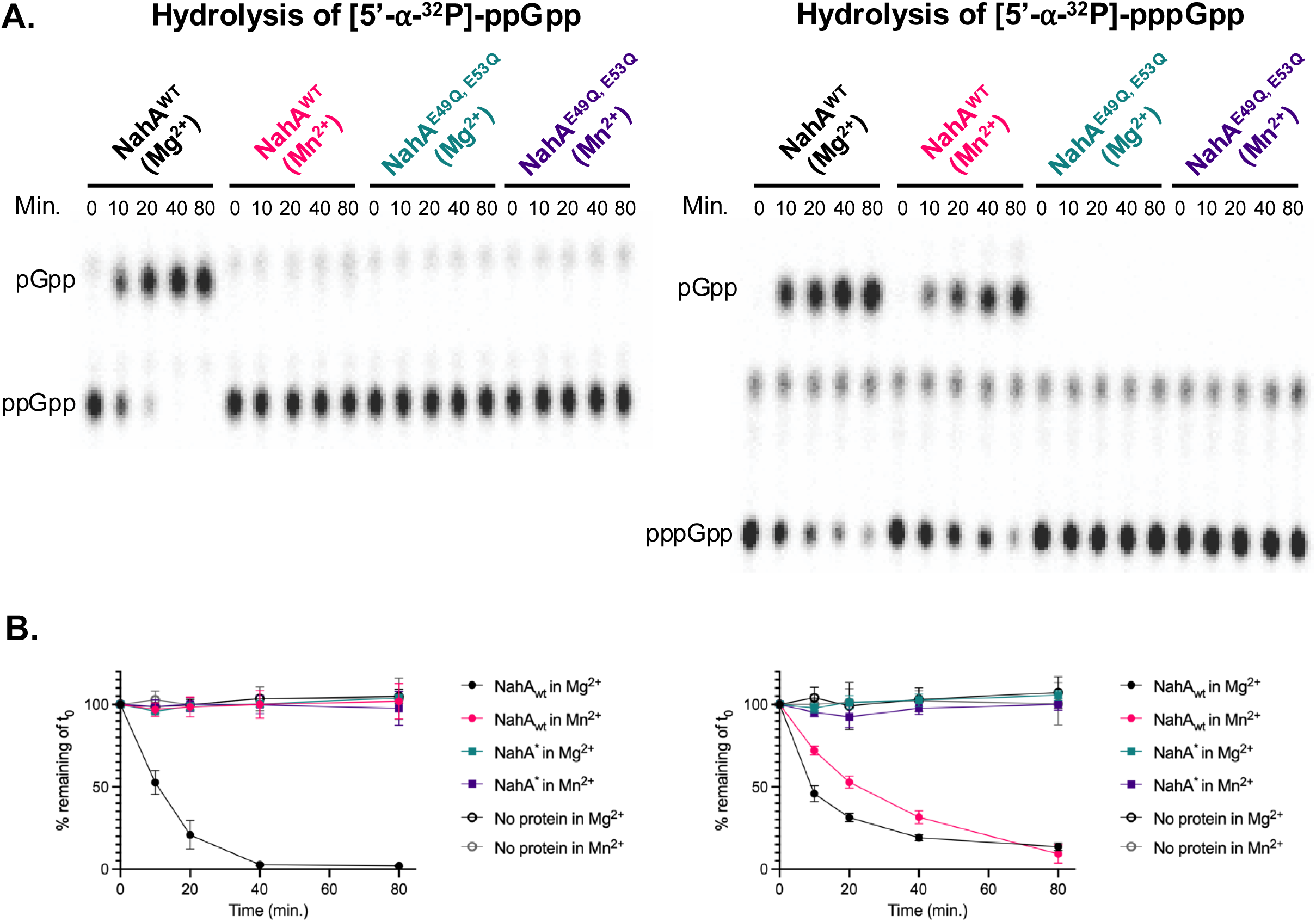
The NuDiX motif is important for (p)ppGpp hydrolysis by NahA. **(A)** *In vitro* hydrolysis activity of ppGpp (left) or pppGpp (right) by NahA into pGpp. Wildtype or catalytically inactive NahA^E49Q, E53Q^ (NahA*) were added to radiolabeled ppGpp or pppGpp and incubated in hydrolysis buffer containing either 1 mM MgCl_2_ or MnCl_2_. **(B)** Quantified hydrolysis from (A).

Additionally, while mRNA pyrophosphohydrolase activity by NahA is strictly manganese dependent *in vitro* (56), we found that hydrolysis of (p)ppGpp was much stronger in magnesium than manganese under the tested conditions; whereas both alarmones are readily hydrolyzed in magnesium, only pppGpp but not ppGpp is hydrolyzed in manganese (Figure 7). In sum, these assays show that the NuDiX motif of NahA is required for (p)ppGpp hydrolysis and, unlike RNA pyrophosphohydrolase that is strictly manganese dependent (56), (p)ppGpp hydrolysis is stronger in magnesium.

## Discussion

Our work using genetic induction of alarmone synthesis yields important insight on the isolated effects of pGpp production, which is an abundant component of the bacterial stress response. While amino acid starvation in wild-type cells leads to a mixed accumulation of all three alarmone species (26), genetic induction of alarmone synthesis in the presence of NahA leads to accumulation of only pGpp and no detectable (p)ppGpp (Figure 2B, Supplemental Figure S1) (39). The difference in the relative abundance of pGpp in these two contexts is likely due the bifunctional synthetase-hydrolase Rel present in wild-type cells and absent in alarmone inducible strains, because Rel hydrolyzes pGpp back to GMP more efficiently than (p)ppGpp to GDP/GTP (26). (Notably, Rel from *Enterococcus faecalis* (15) and *Staphylococcus aureus* (62) have comparable hydrolase activity for all three alarmones.) Therefore, our genetic approach to pGpp and (p)ppGpp production is the first to determine the isolated effects of each alarmone species on translation *in vivo* because it does not invoke exogenous stress that may independently affect translation, such as amino acid starvation.

Using genetic induction of alarmone synthesis, we find that pGpp production inhibits translation *in vivo*, and it does so more rapidly than (p)ppGpp production (Figure 3 and Figure 5). This finding was unexpected since pppGpp and ppGpp are stronger inhibitors of protein synthesis *in vitro* (Figure 1), which is consistent with their higher affinity for translational GTPases (26). Therefore, we propose a model (Figure 8) whereby alarmone production can inhibit translation indirectly *in vivo* by rapidly depleting GTP (Figure 2B), which is sufficient for translation inhibition (Figure 4).

**Figure 8.**
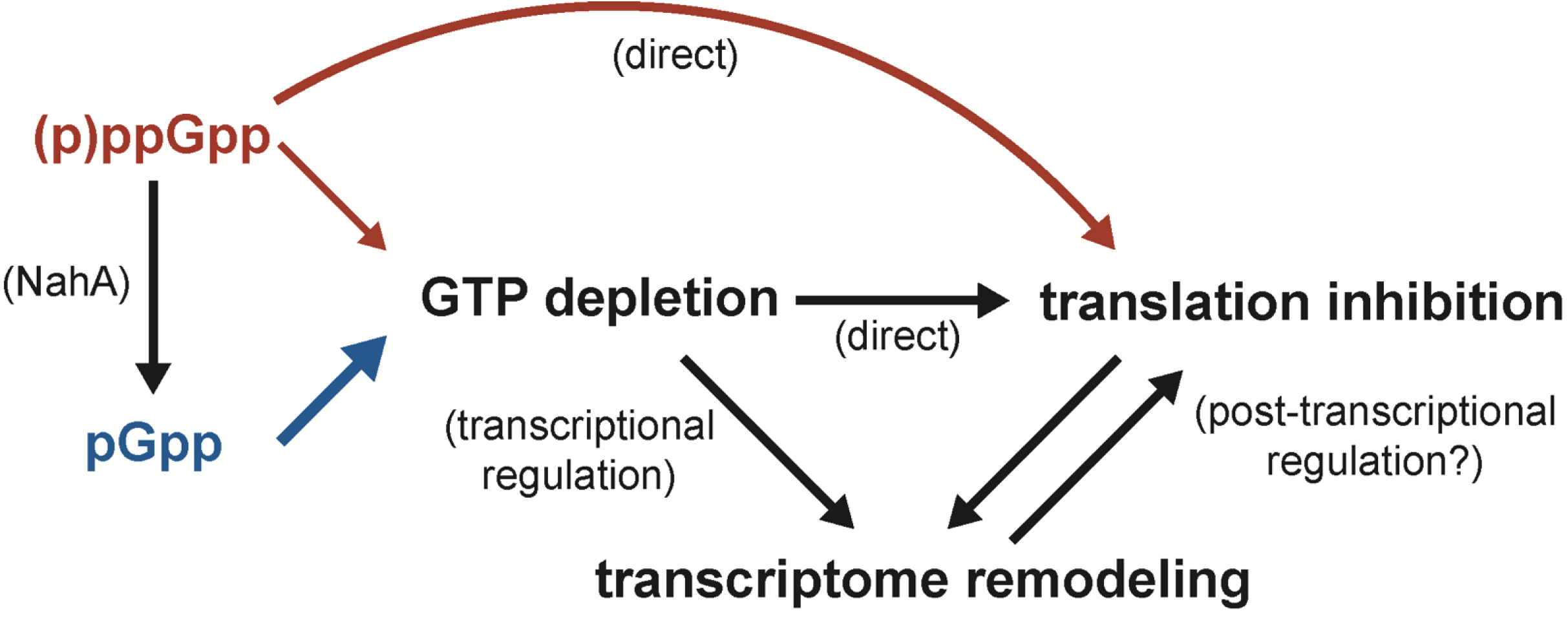
During alarmone production, NahA hydrolyzes (p)ppGpp into pGpp (Figure 2 and Supplemental Figure S1). Starvation-independent production (i.e., genetic induction) of pGpp synthesis depletes GTP modestly more rapidly than (p)ppGpp production (Figure 2), and GTP depletion is sufficient for translation inhibition (Figure 4). GTP depletion may inhibit translation directly by reducing the activity of translational GTPases, or GTP depletion may inhibit translation indirectly as a consequence of transcriptome remodeling. *In vitro*, (p)ppGpp is a stronger inhibitor of protein synthesis than pGpp (Figure 1), due to strong and direct interactions with translational GTPases. *In vivo*, (p)ppGpp production leads to slower translation inhibition than pGpp production (Figure 3 and Figure 5), perhaps due to strong and direct interactions with translational GTPases that may post-transcriptionally affect the transcriptome.

How might ribosomes become disengaged from translation during GTP depletion? Upon GTP depletion by MPA or pGpp induction, we observed rapid polysome run-off (Figure 3 and Figure 4). These results suggest that translation initiation is strongly inhibited and elongation is less affected. One possible mechanism for stronger inhibition of translation initiation than elongation can be explained by the biochemical properties and cellular abundances of translational GTPases. The affinity of EF-Tu for GTP depends on the affinity of the specific aminoacyl-tRNA and the nucleotide exchange factor EF-Ts (29, 63); however, because EF-Tu is the most abundant protein in the cell at approximately 100,000 copies, its GTP-dependent tRNA delivery step is unlikely to become rate-limiting for protein synthesis during GTP depletion. Similarly, EF-G has a low-micromolar K_d_ (i.e., high affinity) for GTP (32, 64) and is highly abundant at approximately 23,000 copies per cell. In contrast, while IF2 has comparable affinity for GTP (65), it is present at only ∼200 copies per cell (66), which may make initiation kinetically sensitive to GTP depletion. Therefore, IF2 is likely to be not only more sensitive to ppGpp in the cell as previously described (28, 30, 32, 33) but also more sensitive to GTP depletion *in vivo*.

(p)ppGpp competitively binds translational GTPases with higher affinity than pGpp (25, 26). Therefore, genetic induction of (p)ppGpp confers strong and direct translation regulation that is exacerbated by GTP depletion. For example, here we observed that (p)ppGpp production leads to fewer disengaged ribosomal subunits after puromycin treatment compared to pGpp production (78% compared to 84%, respectively) (Figure 5, right); this result indicates that (p)ppGpp leads to a larger population of ribosomes that are not compatible with puromycin-mediated termination of the peptide chain. One explanation could be that the puromycin-resistant ribosomes are in a post-transpeptidation but pre-translocation state (i.e., awaiting EF-G), thereby preventing binding of puromycin to the ribosomal A site. Consistent with this observation, IF2 and EF-Tu–but not EF-G–can utilize and hydrolyze pppGpp, which could allow ribosomes to engage with mRNA but not translocate (28, 30, 67, 68). Consistent with this model, at 60 minutes post-alarmone induction where both alarmone inducible strains are equally depleted for GTP (Figure 2B), (p)ppGpp-producing cells have more translationally engaged ribosomes than pGpp-producing cells (Supplemental Figure S2). One potential consequence of this direct translational regulation in the (p)ppGpp inducible strain may be more ribosome protection of mRNA transcripts, which would slow degradation and turnover (69). Supporting this, we found that (p)ppGpp production results in fewer and slower global transcriptomic differences compared to pGpp production (Figure 6, Supplemental Figure S6). Since these differences occur prior to a significant difference in GTP depletion (Figure 2B) and (p)ppGpp does not directly affect global mRNA synthesis in *B. subtilis* (10, 12), we hypothesize that many of these transcriptomic differences between pGpp and (p)ppGpp production reflect differences in post-transcriptional regulation. In total, these observations suggest that, while GTP depletion is sufficient for the bulk of translation inhibition during alarmone production, pppGpp, ppGpp, and pGpp affect translation differently during times when there is comparable GTP depletion.

In addition to faster translation inhibition, we observe that pGpp induction causes a modest but significantly more rapid decrease in GTP levels than (p)ppGpp induction (Figure 2B). There are several potential causes for faster GTP depletion during pGpp production. First, pGpp binds many purine biosynthesis and salvage enzymes with higher affinity than (p)ppGpp (26), and this stronger binding confers stronger inhibition for at least HprT (15, 26), Gmk (15), and XprT (16). Stronger inhibition of Gmk would lead to a greater accumulation of GMP (70), which could increase pGpp production directly through SasA and create a positive feedback loop for Gmk inhibition and GTP depletion. An alternative explanation for faster GTP depletion in the presence of NahA could be that (p)ppGpp hydrolysis makes (p)ppGpp synthesis more thermodynamically favorable; by hydrolyzing the SasA product (p)ppGpp into pGpp, NahA might increase the favorability of pyrophosphorylation of SasA substrates GDP or GTP. Moreover, previous work has shown that RelP/SasA (from *Staphylococcus aureus*) is orthosterically inhibited by high concentrations of (p)ppGpp (71); if GMP is a poor substrate for SasA, then perhaps pGpp is also a poor orthosteric inhibitor and permits greater activity of SasA. This feature of (pp)pGpp regulation would be obscured in a system where alarmone production is not genetically induced because wildtype cells containing the Rel hydrolase would hydrolyze pGpp back to GMP more efficiently than (p)ppGpp to GDP/GTP (26). Although we cannot determine the cause of stronger GTP depletion during pGpp production compared to (p)ppGpp production–nor can we rule out the impact this modest difference might have on translation–the pGpp inducible strain does not accumulate ppGpp or pppGpp (Figure 2B) and therefore lacks strong and direct regulation of translational GTPases.

Because cells producing pGpp and (p)ppGpp exhibit differences in GTP depletion, translation inhibition, and transcriptome remodeling, understanding regulation of (pp)pGpp homeostasis will be critical for deciphering the roles of these nucleotides during stress. One mode of regulation of NahA could be metal coordination, as we establish that (p)ppGpp hydrolysis requires the NuDiX motif, which coordinates metal cofactors (56) (Figure 7). This finding is consistent with previous results showing that EDTA abolishes (p)ppGpp binding to NahA (26). In our hydrolysis assays, the identity of the divalent metal also greatly affects activity (Figure 7), where (p)ppGpp hydrolysis is much stronger in the presence of magnesium than manganese. This observation distinguishes NahA from the other (p)ppGpp hydrolase in wild-type cells, Rel, because hydrolysis by RelA/SpoT homologs (RSHs) are universally manganese-dependent (72–77). Further biochemical characterization of NahA will be needed to determine the range of metal requirements for (p)ppGpp hydrolysis–including the identity of the coordinated metal *in vivo*––and whether this serves to regulate (pp)pGpp homeostasis.

In conclusion, this work implicates all three alarmones in facilitating translation inhibition. We propose that starvation-independent alarmone production inhibits translation largely through GTP depletion. Analogous to our model where GTP depletion strongly inhibits translation initiation by IF2, starvation in eukaryotes (which lack a RelA-like enzyme and therefore cannot produce (p)ppGpp in response to starvation) triggers rapid depletion of NTPs, which causes stronger inhibition of translation initiation than elongation and results in polysome run-off (78, 79). Strong inhibition of the initiation step is a beneficial strategy for cells to maintain functional ribosomes, as it reduces the number of stalled or collided ribosomes that would necessitate alternative modes of ribosome recycling and rescue (80–86). Additionally, some bacteria are proposed to only produce pGpp (40, 41); therefore, direct translational inhibition by (p)ppGpp is not universally required for translation inhibition during alarmone production. Altogether, our work reveals that pGpp production can still facilitate translation inhibition and that all three alarmones have different effects on translation and gene expression.

## Materials and Methods

### Strains and media

Wildtype *B. subtilis* 168 *trpC2* (HAF1) was used as a strain background. All experiments were done in S7 media (87) with modifications (88) at 37 °C, unless specified otherwise.

*sasB*, *sasA*, and *nahA* (if applicable) were sequentially deleted by transforming genomic DNA purified from the respective BKK deletion strains (89) and removing the kanamycin resistance cassette using pDR244 (89). Deletions were PCR confirmed using oligo pairs sasB_chk_F and sasB_chk_R; sasA_chk_F and sasA_chk_R; or KAE12 nahA_F and KAE13 nahA_R, respectively. (pp)pGpp^0^ mutants (KAE55) were generated by transforming Δ*rel*::*kan* genomic DNA from the BKK collection and PCR confirmed using oligos relA_chk_F and relA_chk_R.

To generate a catalytically inactive *nahA* (E49Q E53Q) as previously identified (56), the CRISPR editing plasmid pJW557 (19) (based on pPB41 (90)) was used. The plasmid contained a wildtype *nahA* guide RNA with an appropriately mutated repair template (pJW738). The plasmid was transformed into *B. subtilis*, selected on spectinomycin at 30 °C, and cured at 45 °C. Mutations were screened by Sanger sequencing using oligos KAE12 and KAE13. To obtain this mutation in the (pp)pGpp^0^ background, the same was done in Δ*sasB* Δ*sasA* mutants, before transforming with Δ*rel*::*kan* (resulting in strain KAE213).

For the alarmone inducible mutants, *amyE*::P*_xyl_*-*sasA* was obtained from the Dworkin Lab (JDB4295) (33), and genomic DNA was transformed into Δ*sasB* Δ*sasA* or Δ*sasB* Δ*sasA* Δ*nahA*. Lastly, Δ*rel*::*kan* genomic DNA from the BKK collection was transformed, resulting in pGpp inducible (KAE60) or (p)ppGpp inducible (KAE74) mutants. All alleles were checked by PCR. To mitigate the effects of non-gratuitous induction of *sasA* (due to xylose catabolism and subsequent re-repression of *sasA*), all experiments were done within 1 hour of induction with 1% xylose unless otherwise noted.

To complement *nahA*, plasmid ECE174 was cut with EcoRI and HindIII and Gibson assembled (NEBuilder) with a DNA fragment KAE12 ECE174-nahA-term (Twist). This includes *nahA* with its native promoter and two strong transcriptional terminators. This plasmid was linearized with ScaI and transformed in Δ*sasB* Δ*sasA* Δ*nahA amyE*::P*_xyl_*-*sasA* mutants, before finally transforming with Δ*rel*::kan genomic DNA. *sacA*::*nahA* was confirmed with Sanger sequencing using oligos KAE14-sacA-Sanger-F and KAE15-sacA-Sanger-R.

To delete *hpf*, genomic DNA from Δ*hpf*::*tet* (91) was transformed into Δ*sasB* Δ*sasA* Δ*nahA amyE*::P*_xyl_*-*sasA* and Δ*sasB* Δ*sasA* Δ*nahA* backgrounds and selected on tetracycline. Mutants were then transformed with Δ*rel*::kan genomic DNA. *hpf* deletion was PCR confirmed using oligos HF15 and HF17.

Plasmid for His-SUMO-NahA^E49Q, E53Q^ overexpression (pJW740) was made in a similar way as the previously published pJW739 plasmid (26), except that the *nahA^E49Q, E53Q^* sequence was PCR amplified using primers oJW3519/oJW3520 from gDNA of *nahA^E49Q, E53Q^*strain made by CRISPR (made with pJW738, see above).

### RNA-sequencing during alarmone induction

For RNA-sequencing before and after alarmone induction, cultures were repeatedly sampled and extracted for total RNA. Briefly, 3 mL of culture was pelleted, resuspended in EDTA, and lysed by incubating with ∼1 mg/mL lysozyme at 37 °C for 10 min. 1 mL of Trizol was added and incubated for 5 minutes at room temperature. 200 µL of chloroform was added, inverted to mix for 15 s, and incubated at room temperature for 10 min. Samples were then centrifuged for 10 min. at 4 °C at maximum speed. The upper phase was transferred to a new tube. 500 µL isopropanol was added and inverted 6-8 times to mix. Samples were precipitated for 10 min. at room temperature and then incubated at -80 °C overnight. The next day, RNA was pelleted by centrifuging for 15 min. at 4 °C at maximum speed. Pellets were washed twice with 1 mL ice-cold 75% ethanol. Pellets were dried for 1 minute at room temperature, before gently resuspending in water.

Total RNA was sent to SeqCenter (Pittsburgh, PA) for library preparation, rRNA depletion, and paired-end RNA-sequencings (NovaSeq X Plus, 20M reads). Sequencing data was mapped to NC_000964.3 with Bowtie2 (2.5.1) and quantified with FeatureCounts. Differential expression was determined using DESeq2 (92). Because cultures (i.e., samples) were repeatedly measured over time, we used the following design to compare differential expression within strains and across strains over time,

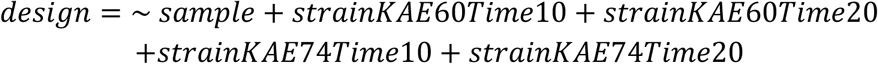

As an example, to obtain log2FoldChanges and adjusted p-values to compare 10 minutes post-induction to 0 minutes for just strain KAE60, we used,

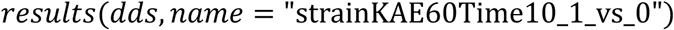

To obtain log2FoldChanges and adjust p-values that reflect strain-dependent effects on expression over time, we used,

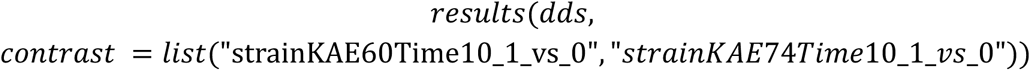

### Wildtype vs. *nahA*^E49Q,^ *^E53Q^* transcriptome analysis and rifampicin-sequencing (rif-seq)

For analysis of the steady-state transcriptome, wildtype and *nahA*^E49Q, E53Q^ were harvested and RNA was extracted essentially as described (93, 94). Total RNA (N = 2) was sent to SeqCenter (Pittsburgh, PA) for library preparation and paired-end RNA-sequencing (NovaSeq X Plus, 20M reads). Sequencing data was mapped to NC_000964.3 with Bowtie2 (2.5.1) and quantified with FeatureCounts. Differential expression was determined using DESeq2.

To approximate mRNA half-lives, rifampicin-sequencing (rif-seq) was performed. Cultures were repeatedly harvested, RNA extracted, and sequenced pre-rifampicin treatment (*t =* 0), 2 minutes post-, and 5 minutes post-rifampicin treatment (150 µg/mL). Sequences were mapped with Bowtie2 and quantified with FeatureCounts. Counts for each gene were first divided by the length of each gene to obtain counts per base. Lowly expressed genes that had a counts per base value less than *sasA*, *sasB,* and *rel* (which were deleted) were omitted. Then, these values were normalized by dividing by the sum of the counts mapping to *ssrA* and *rnpB* within each sample, which are known abundant and stabile RNAs, to obtain normalized counts per base. To mitigate the confounding effect that runoff of elongating RNA polymerase has on RNA levels post-rifampicin treatment(95–97), only genes within a monocistronic or dicistronic operon were selected. Next, the percent remaining normalized counts per base was calculated for each gene by dividing the *ssrA/rnpB*-normalized counts per base at *t* = 0. A linear model was fit to the log-transformed percent remaining normalized counts per base, where the y-intercept was forced to log(100). Lastly, half-life was calculated by dividing -log(2) by the slope of the linear model. To test for statistical significant differences in a gene’s half-life across the two strains, a Student’s T-test was performed.

### Polysome profiling

Strains were grown overnight at room temperature in 4 mL of S7 media and sub-cultured to OD 0.05 in 4 mL. Starter cultures were grown to OD ∼1.0 at 37 °C, and then sub-cultured to OD 0.01 in 250 mL in a 2 L baffled flask. Cultures were incubated at 37 °C with shaking until they reached an OD 0.4-0.7. This marked t = 0 (pre-induced). 35-40 mL of cells were collected by centrifugation at 10000 rpm for 5 minutes (Beckman Coulter Avanti J-15R, rotor JA-10.100), the supernatant poured off, and cell pellet stored at -80 °C. Xylose was quickly added to a final concentration of 1% to induce *sasA* expression, or MPA was added to a final concentration of 80 µg/mL to inhibit GuaB. Cells were harvested at the time-points specified in the figures.

Cell pellets were washed and resuspended with cold gradient buffer (20 mM Tris, 60 mM NH_4_Cl_2_, 7.5 mM MgAoc, 6 mM 2-mercaptoethanol, 0.5 mM EDTA). lysed by bead beating 5 times on maximum speed for 30 seconds (Benchmark Scientific BeadBug 6), with 2 minutes on ice in between. For the ribosome dissociation experiments, puromycin was added to a final concentration of 100 µg/mL just prior to lysis. S30 extracts were collected by centrifuging the samples at 21300 rcf for 20 minutes (Eppendorf Centrifuge 5425 R).

Sucrose density gradients were prepared using 10% and 40% sucrose dissolved in gradient buffer and using a Biocomp Gradient Station (Biocomp Instruments). For the ribosome dissociation experiments, Ribosome concentration in the S30 extracts were approximated by Abs_260_ (NanoDrop), and equal quantities were loaded onto the gradients (typically 250 µg). Gradients were centrifuged for 3 hours at 30,000 rpm in an SW-41Ti rotor. Gradients were read and collected using a Biocomp Gradient Station. Profiles were quantified using QuAPPro (98).

### Western analysis

25 µL of each gradient fraction was mixed with 75 µL 4X loading buffer. Mixtures were heated at 90 °C for 5 minutes and cooled on ice. 20 µL were separated by 12% SDS-PAGE. For each strain, the region of the gel corresponding to the size of HPF was cut and transferred to the same membrane (BioRad Immun-Blot PVDF Membrane), and the same was done for EF-Tu. (I.e., Time-points for each strain were transferred to the same membrane.) Membranes were blocked for 30 minutes with 3% bovine serum albumin (BSA) in PBS-T before adding polyclonal antibodies raised against HPF (91) or EF-Tu (99) overnight at 4 °C. Secondary antibody (anti-rabbit) conjugated to HRP was added for detection. Membranes were washed 3 times with PBS-T and developed with ECL reagent and imaged on ChemiDoc MP (BioRad). Bands were quantified using Image Lab software (BioRad).

### HPG labeling (total protein synthesis assay)

Concurrent to harvesting cells for polysome profiling, 0.5 mL of cells were transferred to culture tubes at labeled with *L*-homoproparyglglycine (Vector Laboratories) at a final concentration of 100 µM to assay total protein synthesis. Cells were labeled for 15 minutes, pelleted, and fixed in 4% formaldehyde (in PBS) for 15 minutes. After washing in 3% BSA in PBS, cells were permeabilized with 0.5% Triton X-100 in PBS for 15 minutes. After washing, final Click reactions using the AlexaFluor 488 and “Click-iT Cocktail” according to manufacturer protocols (Invitrogen). Fluorescence microscopy was done using the Nikon ECLIPSE Ni-E microscope with GFP-FITC filter cube, with a 20 ms exposure time for GFP. Phase contrast and fluorescence channels were combined, background-subtracted, and quantified using ImageJ. For the full time-course assay in Figure 3A, cells were removed from 25 mL cultures (instead of 250 mL cultures that were also used for polysome profiling).

### (pp)pGpp nucleotides preparation and purification

[5′-α-^32^P]-radiolabeled nucleotides pGpp, ppGpp and pppGpp were synthesized and purified as previously described (26). A similar approach using non-radiolabeled GTP was used to make non-radiolabeled alarmones.

### Overexpression and purification of NahA and NahA^E49Q, E53Q^

For proteins overexpression, we used *E. coli* BL21(DE3) lacI^q^ cells transformed either with His-SUMO-NahA vector (pJW739) or the His-SUMO-NahA^E49Q, E53Q^ vector (pJW740). His-tagged NahA and NahA^E49Q, E53Q^ were purified by Ni-NTA column, followed by SUMO protease cleavage to remove the tag as done previously (26).

### *In vitro* (p)ppGpp hydrolysis assays

Kinetic assays of pppGpp and ppGpp hydrolysis were adapted from a previous study (26). Briefly, 100 nM of purified NahA protein, the catalytic mutant NahA^E49Q, E53Q^, or a corresponding volume of protein storage buffer (50 mM Tris-HCl pH8, 500 mM NaCl, 5% glycerol v/v, and 1 mM β-mercaptoethanol) were used for hydrolysis assays and controls, respectively. Assays were conducted at 37 °C in reaction buffer containing either manganese or magnesium (40 mM Tris-HCl pH7.5, 10 mM NaCl supplemented with 1 mM metal chloride) in presence of 0.1 nM [5′-α-^32^P]-pppGpp or [5′-α-^32^P]-ppGpp and 100 μM of the corresponding cold nucleotide. At each indicated timepoint, samples were collected by mixing 10 μL of the reaction mix with 10 μL ice-cold 0.8 M formic acid to stop the reaction. We resolved samples by thin-layer chromatography on PEI-cellulose plates with 1.5 M KH_2_PO_4_ (pH 3.4) (MilliporeSigma), exposed plates to a phosphor screen, and scanned the screen on a Typhoon FLA9000 scanner (GE Healthcare). Nucleotides levels were quantified on Fiji (NIH).

### *In vitro* translation

*In vitro* translation was performed according to manufacturer instructions (NEB PURExpress), with reactions scaled down to 5 µL and incubated for 2 hrs. DHFR control plasmid was added to a final concentration of ∼6 ng/µL. 2 µL of 4X SDS loading buffer was added to the samples, heated at 90 °C for 3 min., and crash-cooled on ice. 5 µL was separated by 12% SDS-PAGE and stained with Coomassie stain. Bands were quantified using Image Lab software (BioRad). As a loading control, densitometric values were normalized to the most-intense band in the gel, which we speculate is EF-Tu based on molecular size. Because the kit contains a protein of similar size to DHFR, we also subtracted the EF-Tu normalized signal from a reaction containing no DHFR control plasmid. The background and no-templated subtracted, EF-Tu normalized values at each alarmone concentration was divided by the 0 alarmone value to generate the fold-change values reported in Figure 2.

### *In vivo* ^32^P labeling, nucleotide extraction, and thin-layer chromatography (TLC)

0.5 or 1 mL of cultures grown in low-phosphate S7 media (0.1X phosphate) were labeled with 25 or 50 µL of ^32^P-orthophosphate by incubating from OD ∼0.02 to OD ∼0.2. 100 µL cells were incubated with 20 µL 2 M formic acid on ice for at least 20 minutes. Samples were centrifuged at 4 °C at 14000 rcf for at least 15 minutes. The top 50 µL was transferred to a new tube. 2 µL was spotted onto PEI cellulose TLC plates (Millipore) and resolved in 1.5 M or 0.85 M KH_2_PO_4_ (pH 3.4) to resolve (p)ppGpp or GTP respectively. TLC plates were exposed to phosphor screens overnight and scanned on Typhoon imager. Spots were quantified using Image Lab software (BioRad). Signals were normalized to the pre-treatment OD and reported as fold-change from untreated samples

**Table 1.**
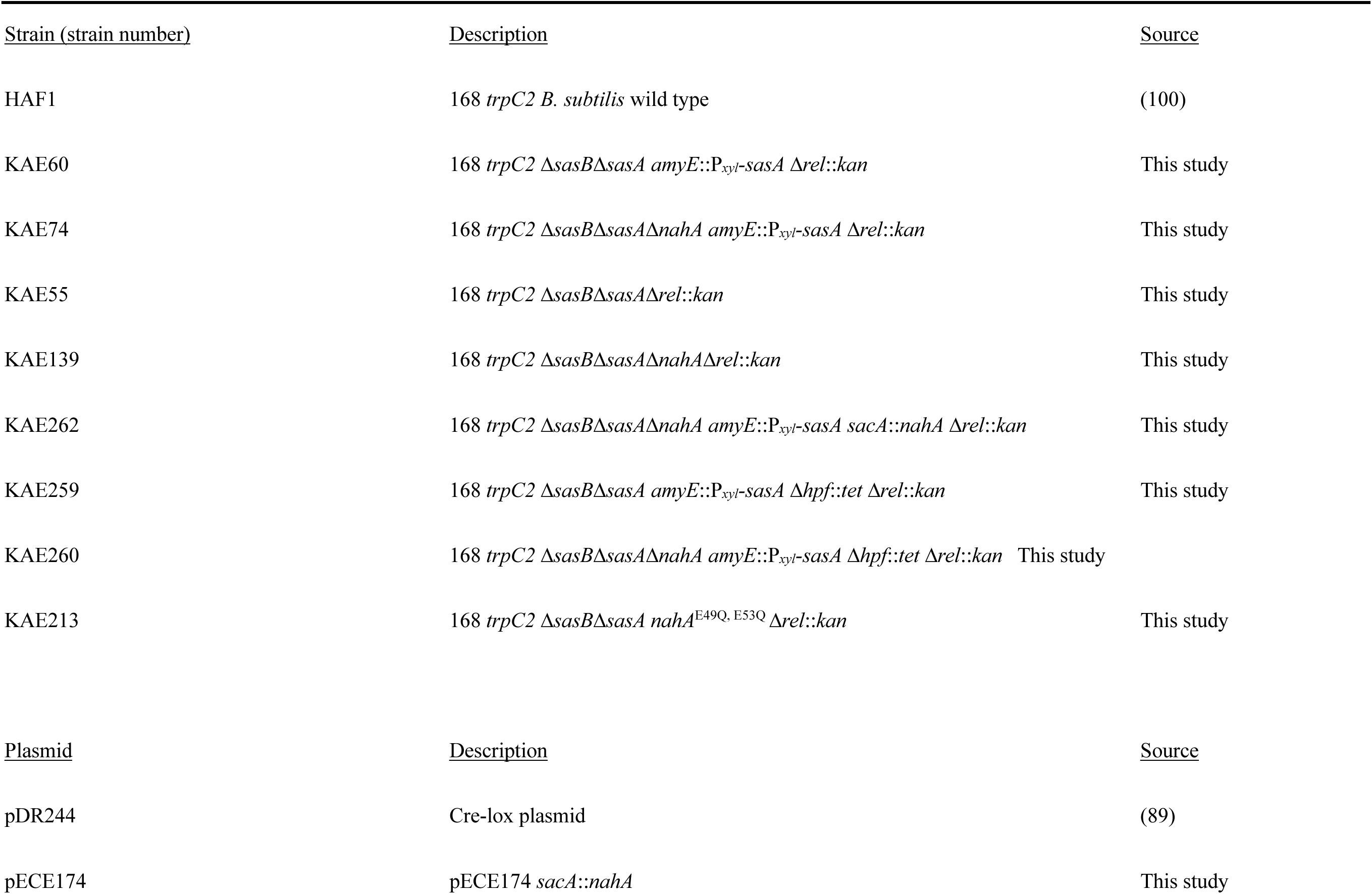
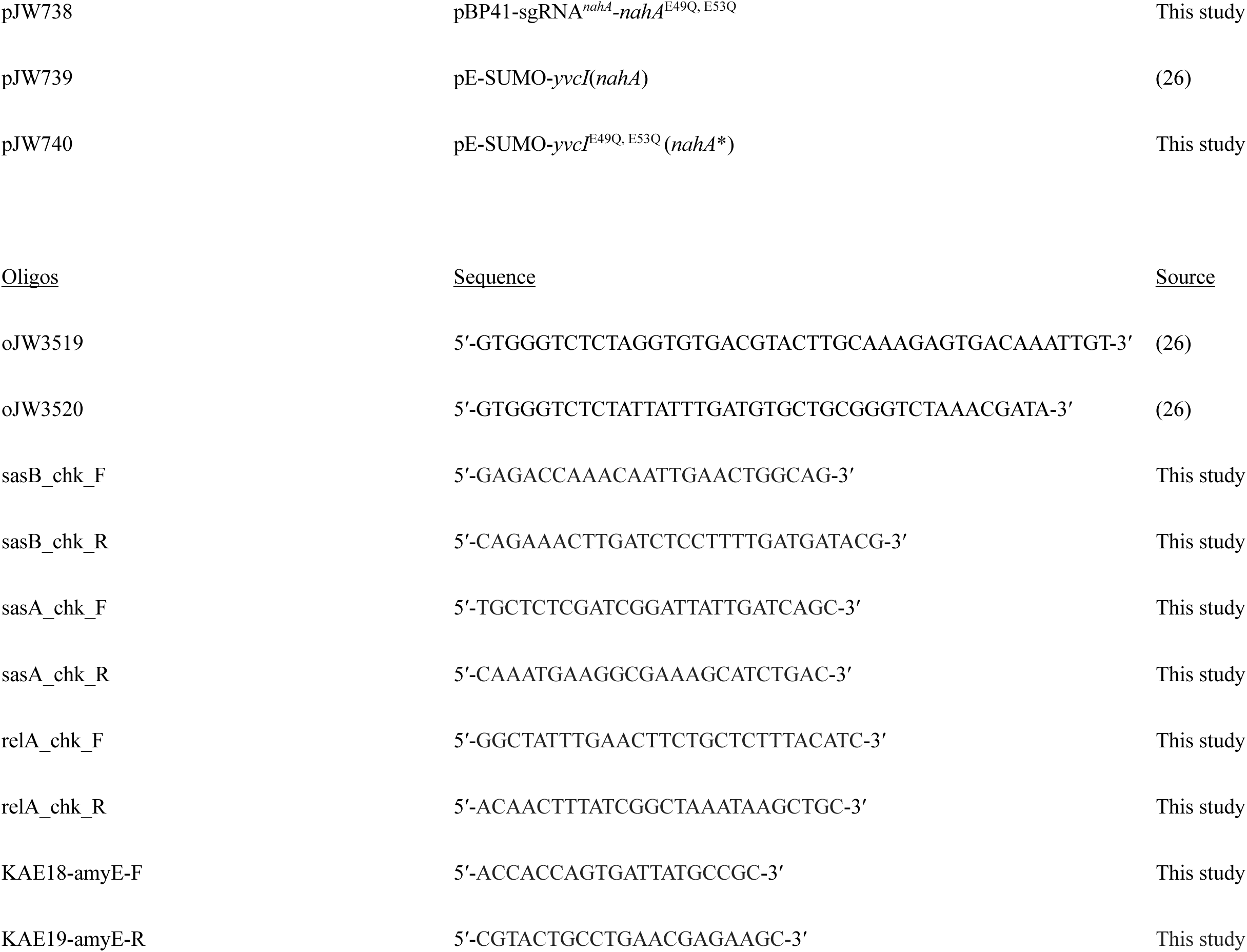
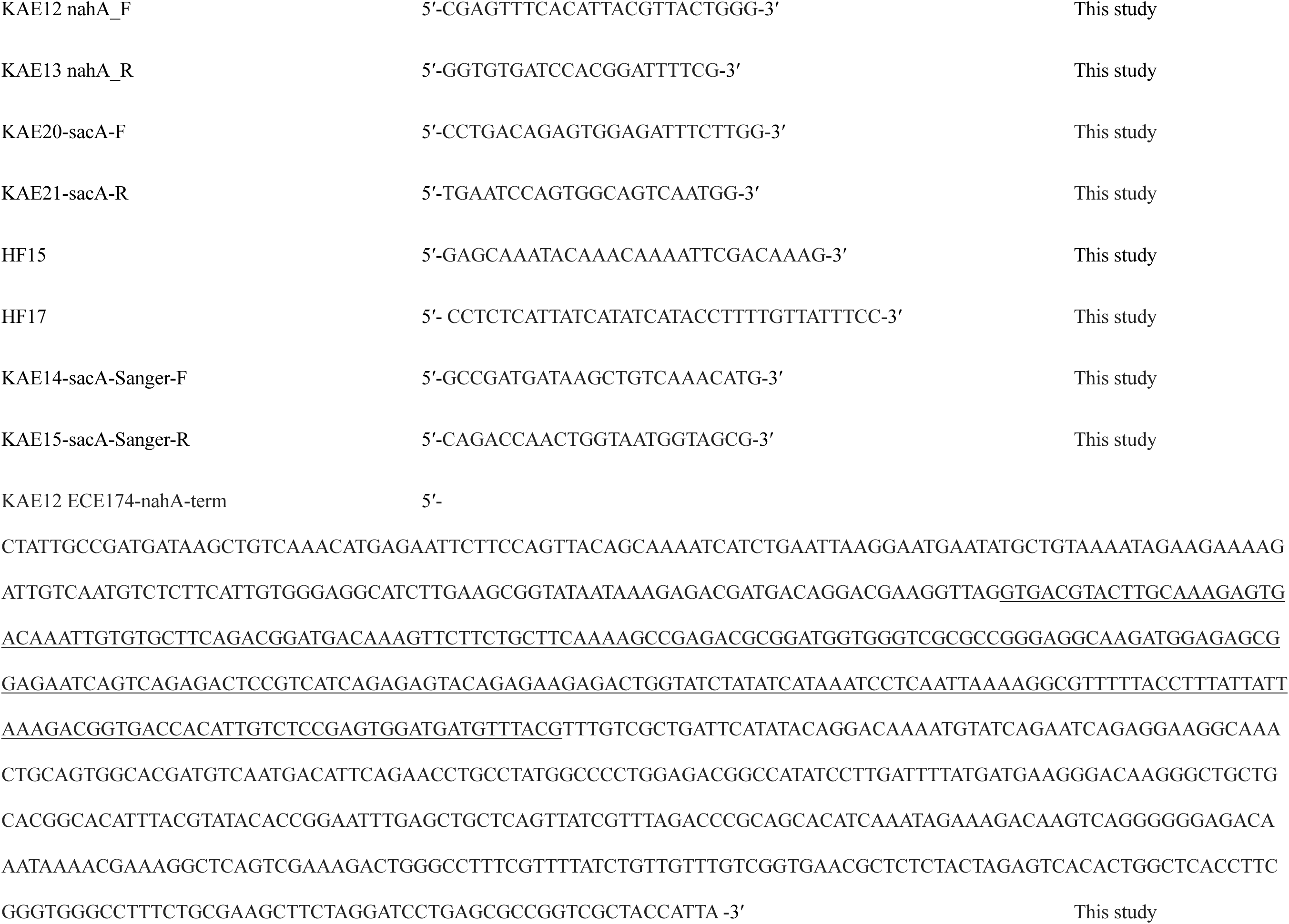
Strains, plasmids and primers.

## Acknowledgements

We would like to thank the Cornell Statistical Consulting Unit for assisting with statistical and differential expression analyses; Jin Yang for creating plasmids pJW738, pJW739, and pJW740; Katrina Callan for assistance with western blotting; Cassidy Prince for assistance with Bowtie2 and featureCounts for RNA-seq; Hannah Leblanc for Rif-seq technical suggestions; Katrina Callan, Cassidy Prince, and Daniel Tetreault for assistance with harvesting cultures for Rif-seq; Danny Fung for TLC analysis advice and suggestions; Augustus Pendleton for coding assistance; and John Helmann for discussions and critical feedback. This work is funded by NIH R35GM147049 to HAF and NIH R35GM127088 to JDW.

---

**Supplemental Figure S1.**
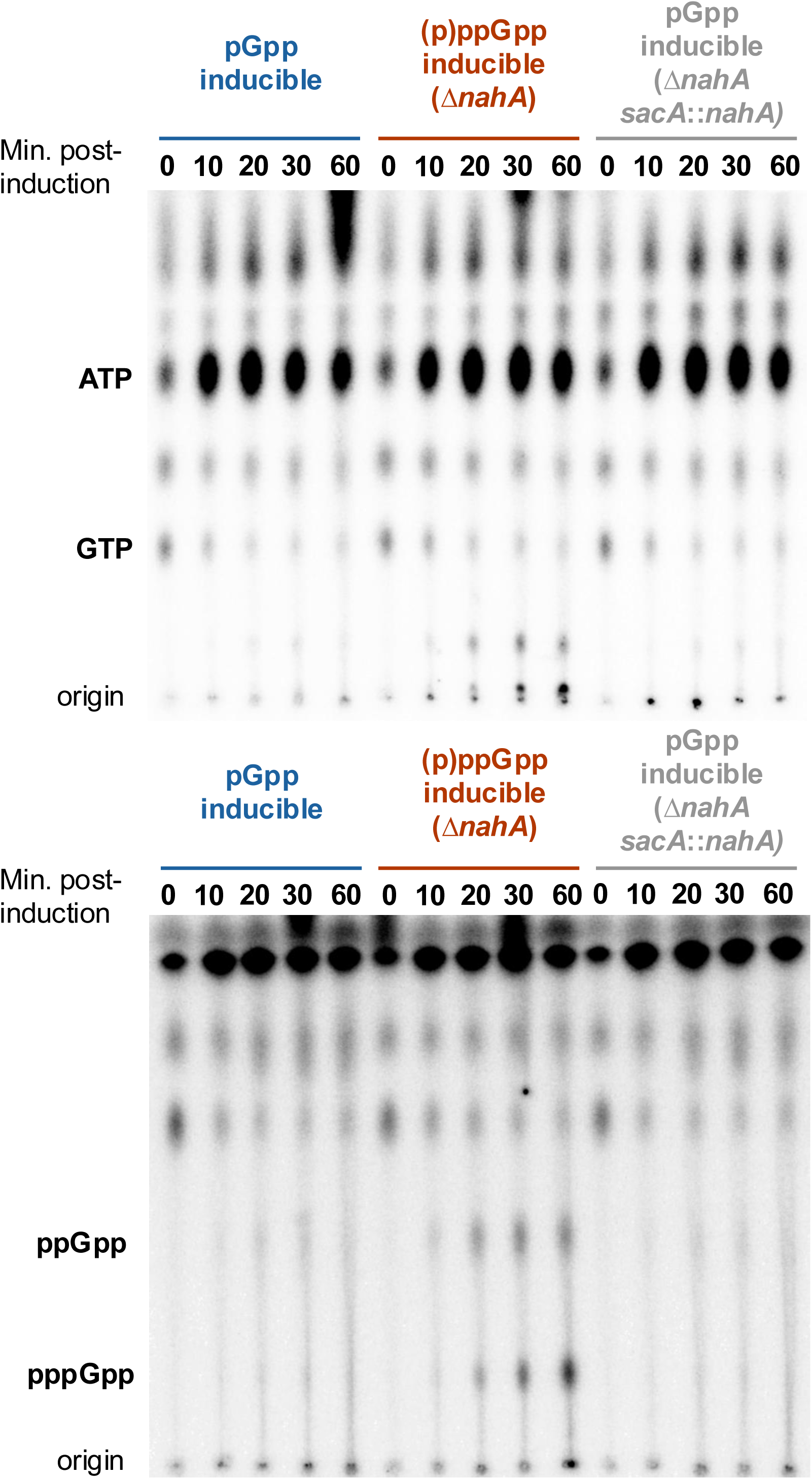
Representative TLC plates of *in vivo* ^32^P-labeled nucleotides in pGpp inducible and (p)ppGpp inducible (Δ*nahA*) strains, with *nahA* complementation (Δ*nahA*, *sacA*::*nahA*). Plates show ATP and GTP (top) or ppGpp and pppGpp (bottom), resolved in 0.85 M or 1.5 M KH_2_PO_4_ (pH 3.4) respectively.

**Supplemental Figure S2.**
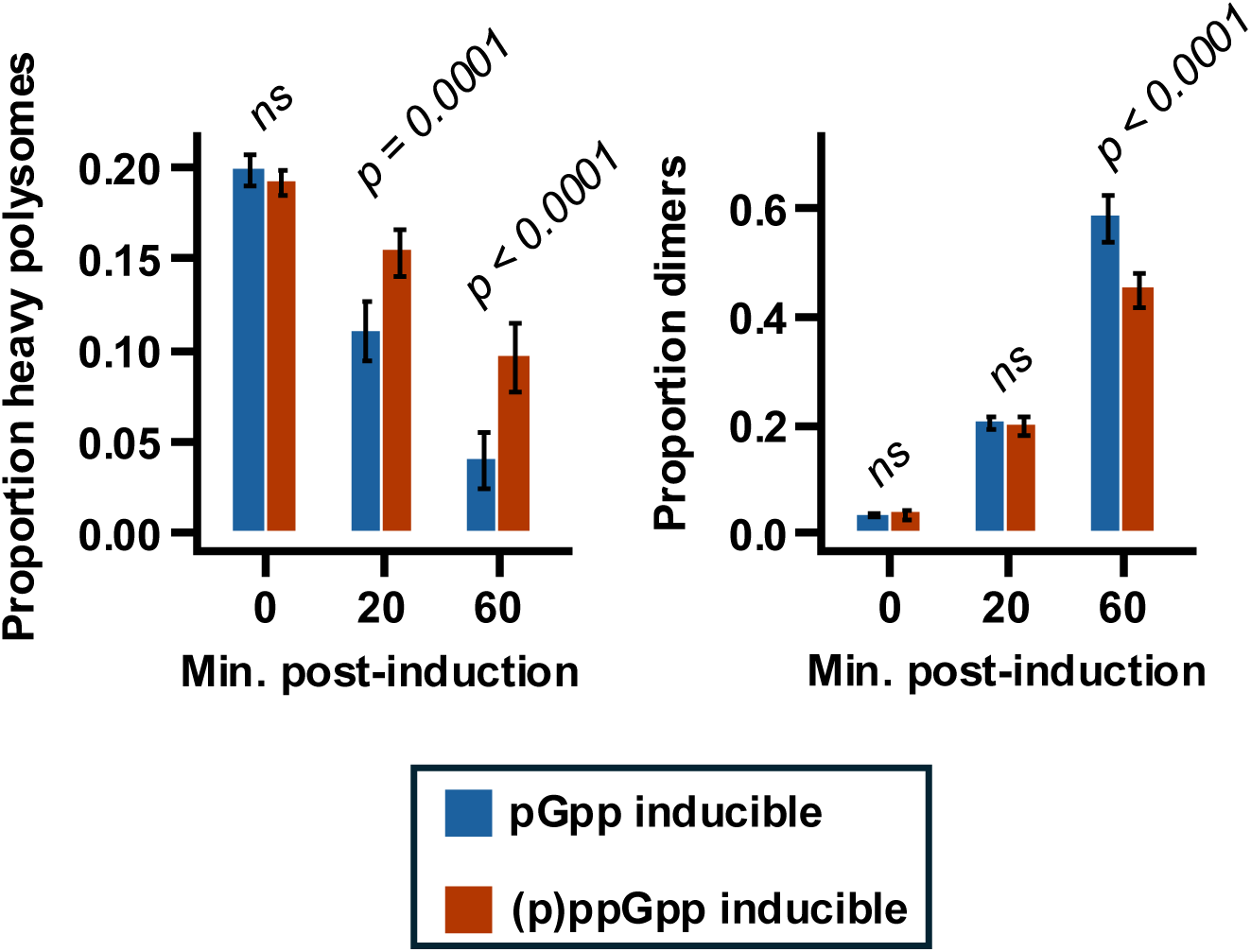
pGpp induction decreases heavy polysomes and increases hibernating ribosome dimers faster than (p)ppGpp induction. (Area under the heavy polysomes and dimer peaks was determined from profiles in Figure 4, Supplemental Figure S5, and one additional replicate not shown). Values represent means (*n =* 3, +/- S.D.). A linear mixed-effects model was used for each figure, where time post-induction and strain are categorical fixed effects and replicate a random effect; *P*-values represent statistical differences between the strains at each time.

**Supplemental Figure S3.**
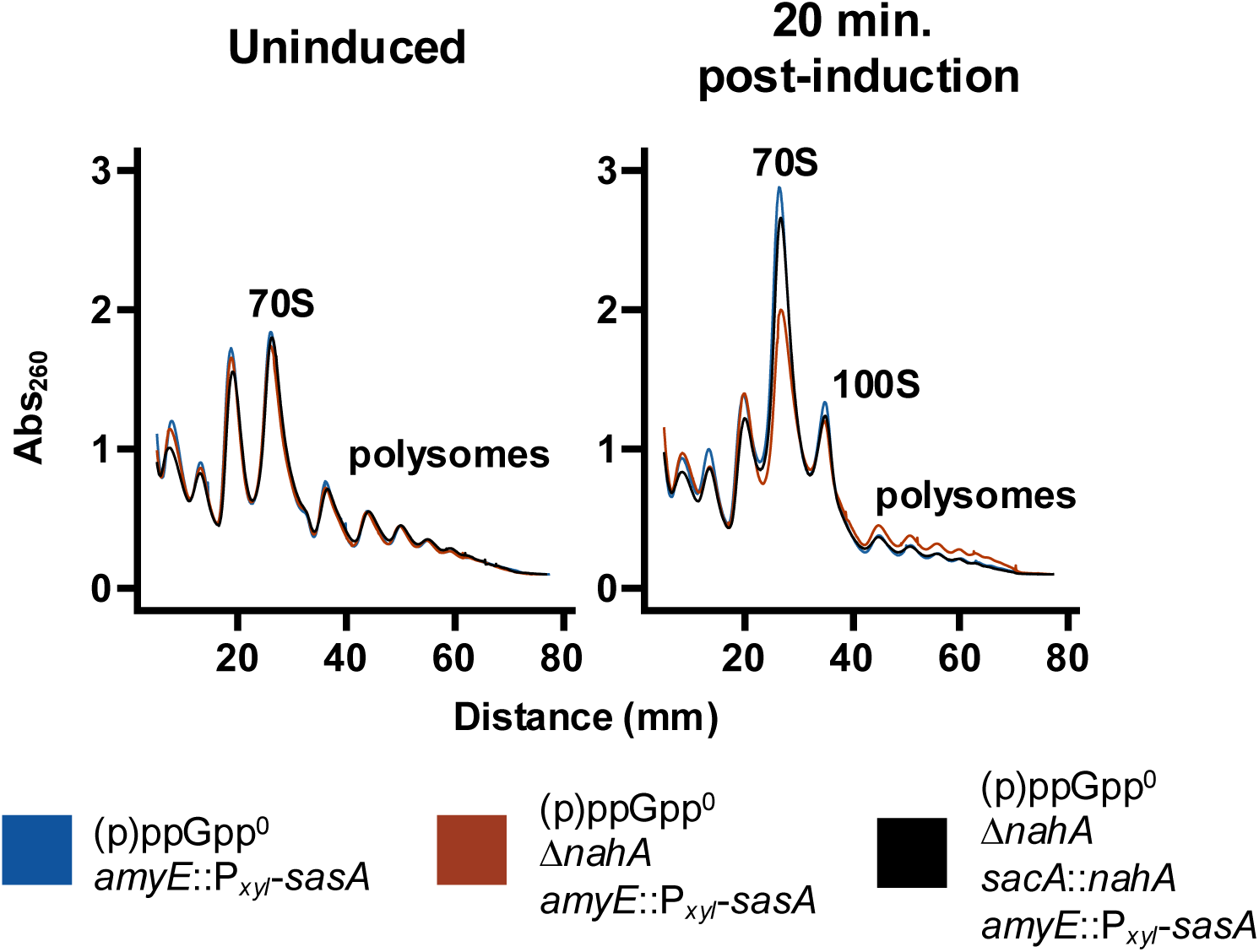
*nahA* complementation restores increased 70S ribosomes and reduced polysomes 20 minutes post-induction of alarmone production.

**Supplemental Figure S4.**
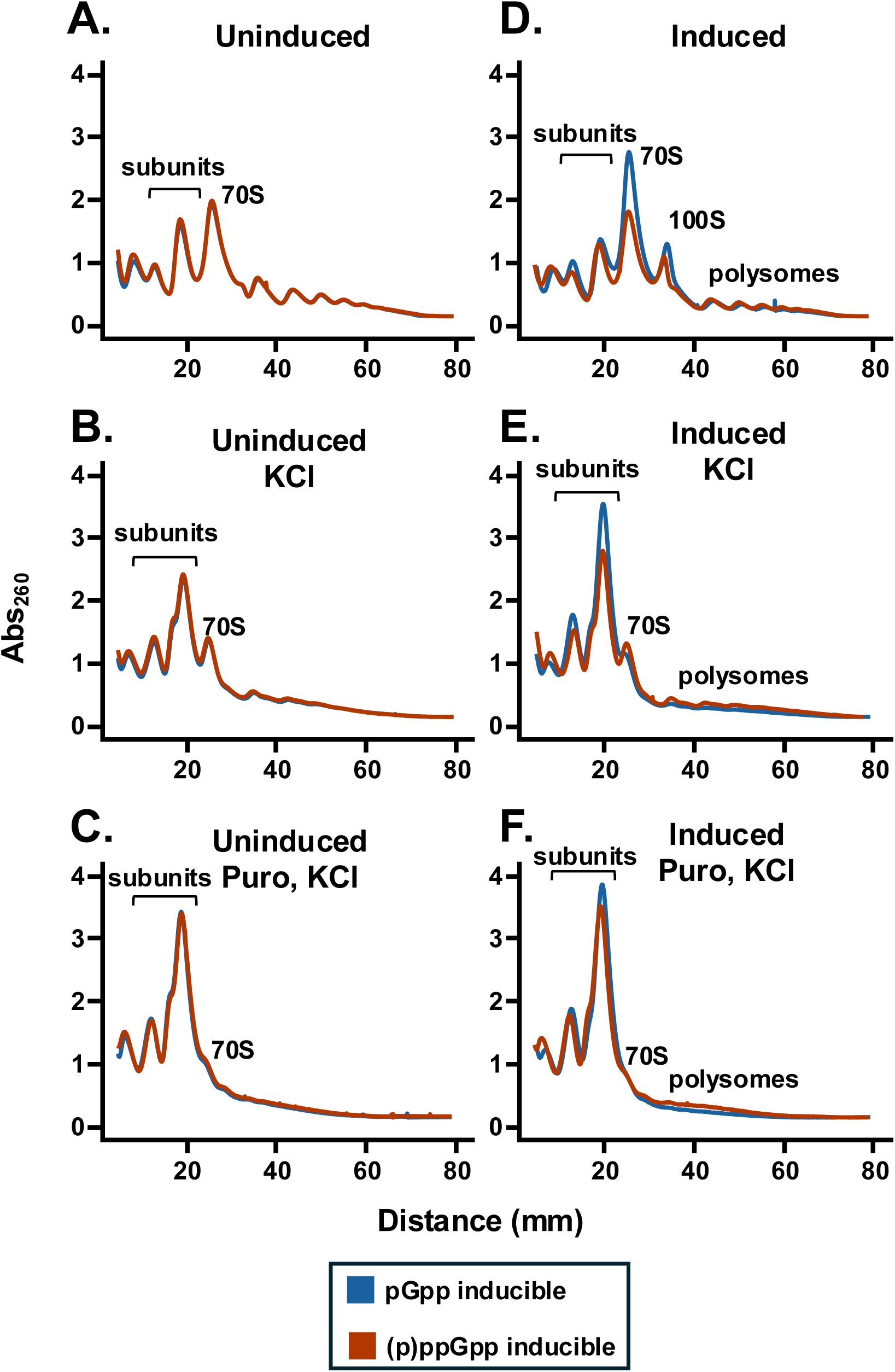
Representative polysome profiles of pGpp and (p)ppGpp inducible mutants containing HPF, quantified in Figure 5.

**Supplemental Figure S5.**
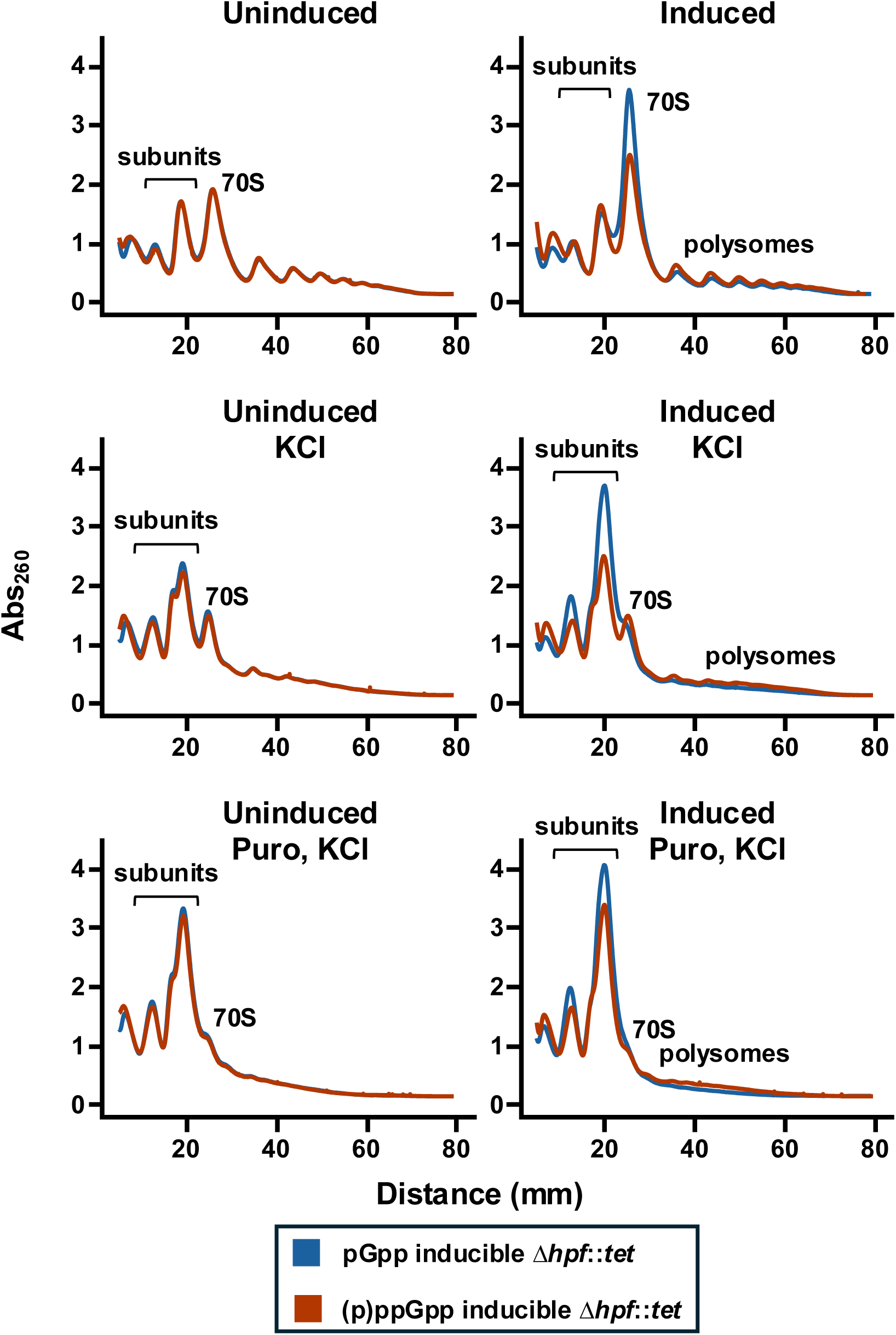
Representative polysome profiles of pGpp and (p)ppGpp inducible mutants lacking HPF (Δ*hpf::tet*), quantified in Figure 5.

**Supplemental Figure S6.**
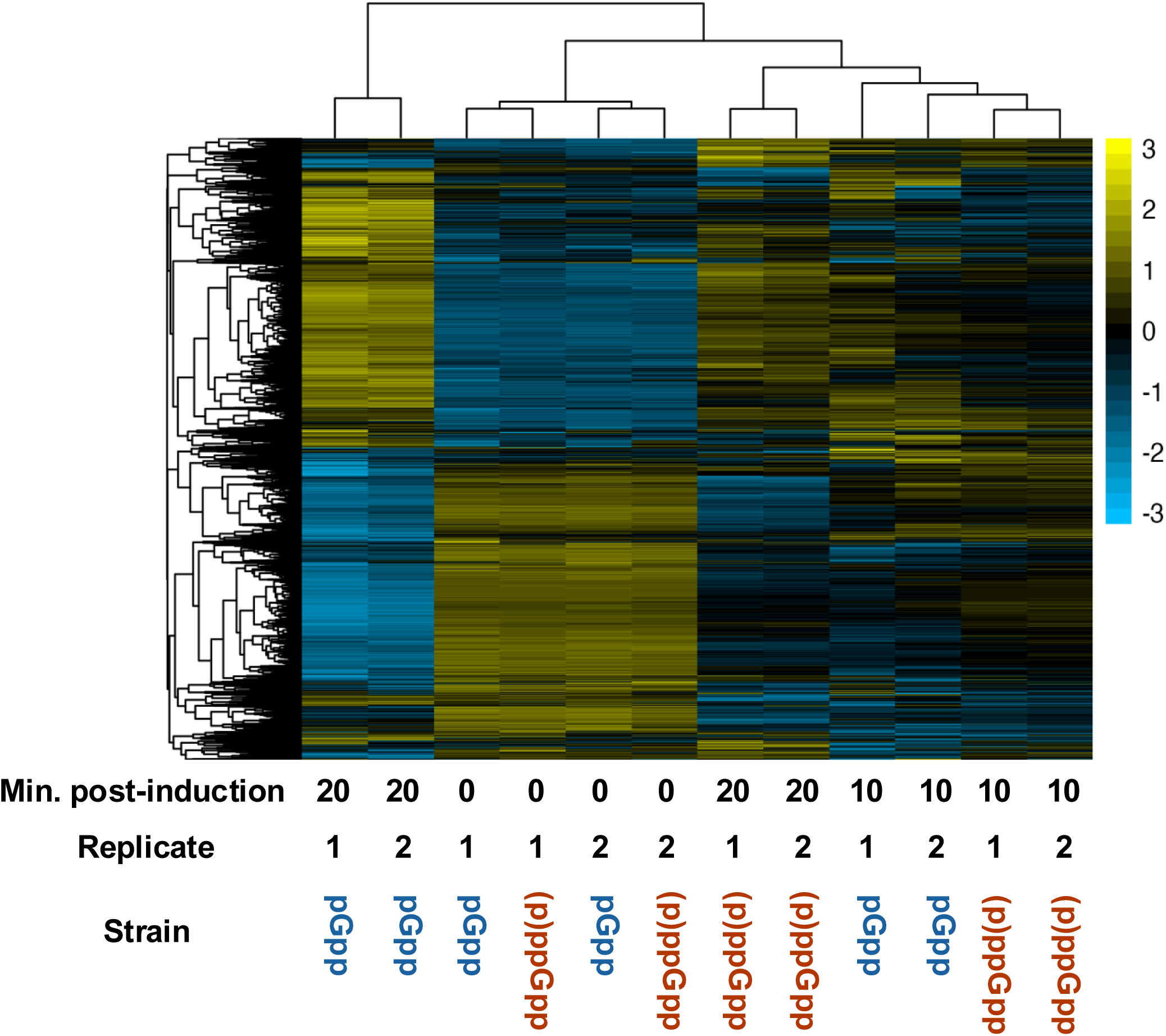
Hierarchical clustering of RNA-seq data, where columns are samples and rows are genes. A hierarchical clustering map with readable gene names is in Supplemental Dataset S2.

**Supplemental Figure S7.**
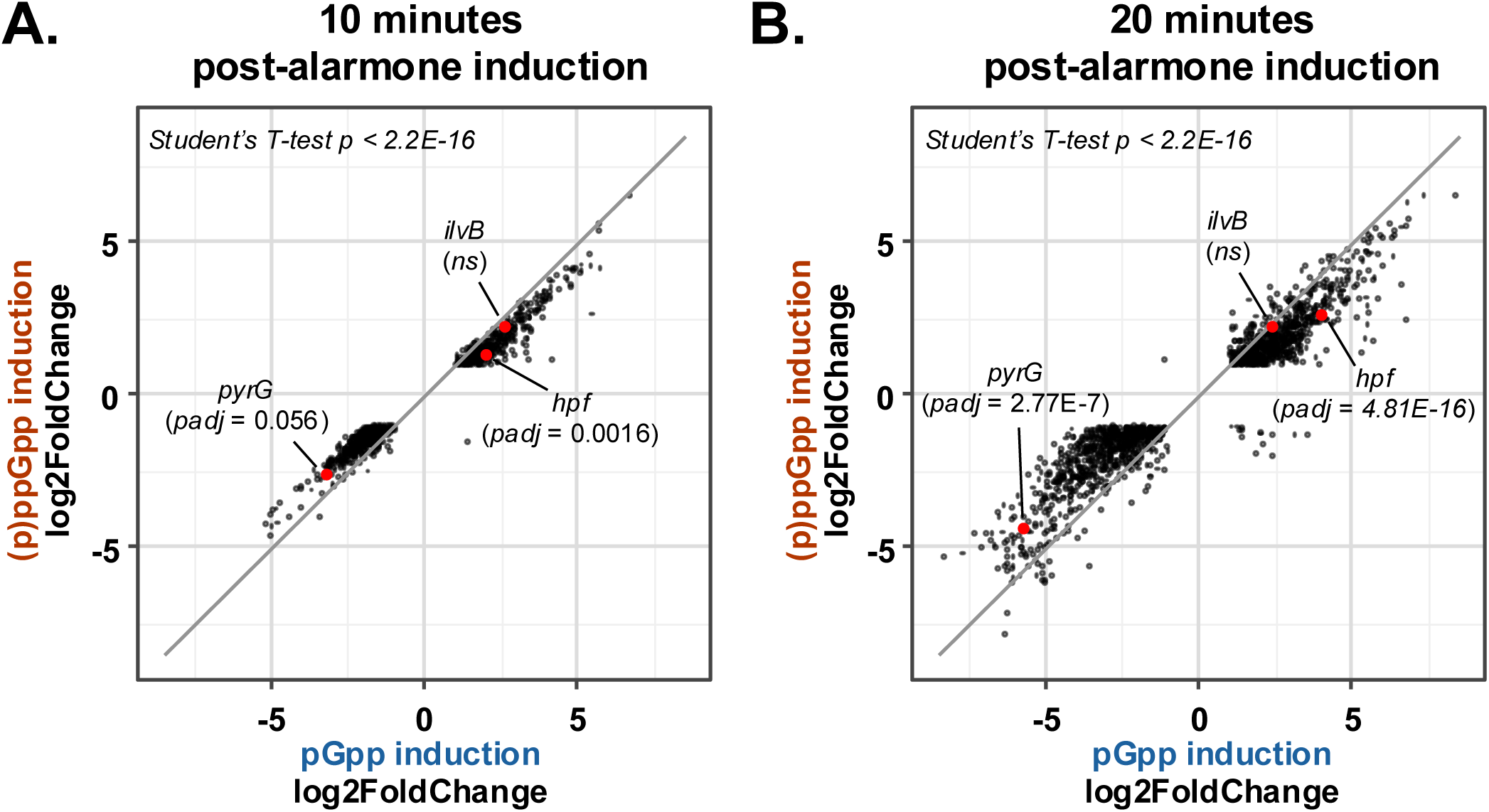
Among the genes that are differentially expressed during pGpp or (p)ppGpp induction, they are differentially expressed more during pGpp production. Points represent genes that are differentially expressed after pGpp or (p)ppGpp induction. X-coordinates represent differential expression pGpp induction relative to pre-induction, and y-coordinates represent differential expression in (p)ppGpp induction relative to pre-induction. The gray lines represent the expected trendline of the data if all genes had equal changes in expression during pGpp and (p)ppGpp induction. Transforming log2FoldChanges to the absolute values and comparing means between pGpp and (p)ppGpp induction using a paired T-test gives *P* < 2.2E-16 at both 10 minutes and 20 minutes post-induction. *pyrG*, *hpf,* and *ilvB* are examples of genes that are more strongly repressed, activated, or unchanged by pGpp induction compared to (p)ppGpp induction; *padj* represents the strain-dependent statistical differences of those genes, relative to t = 0 min.

**Supplemental Figure S8.**
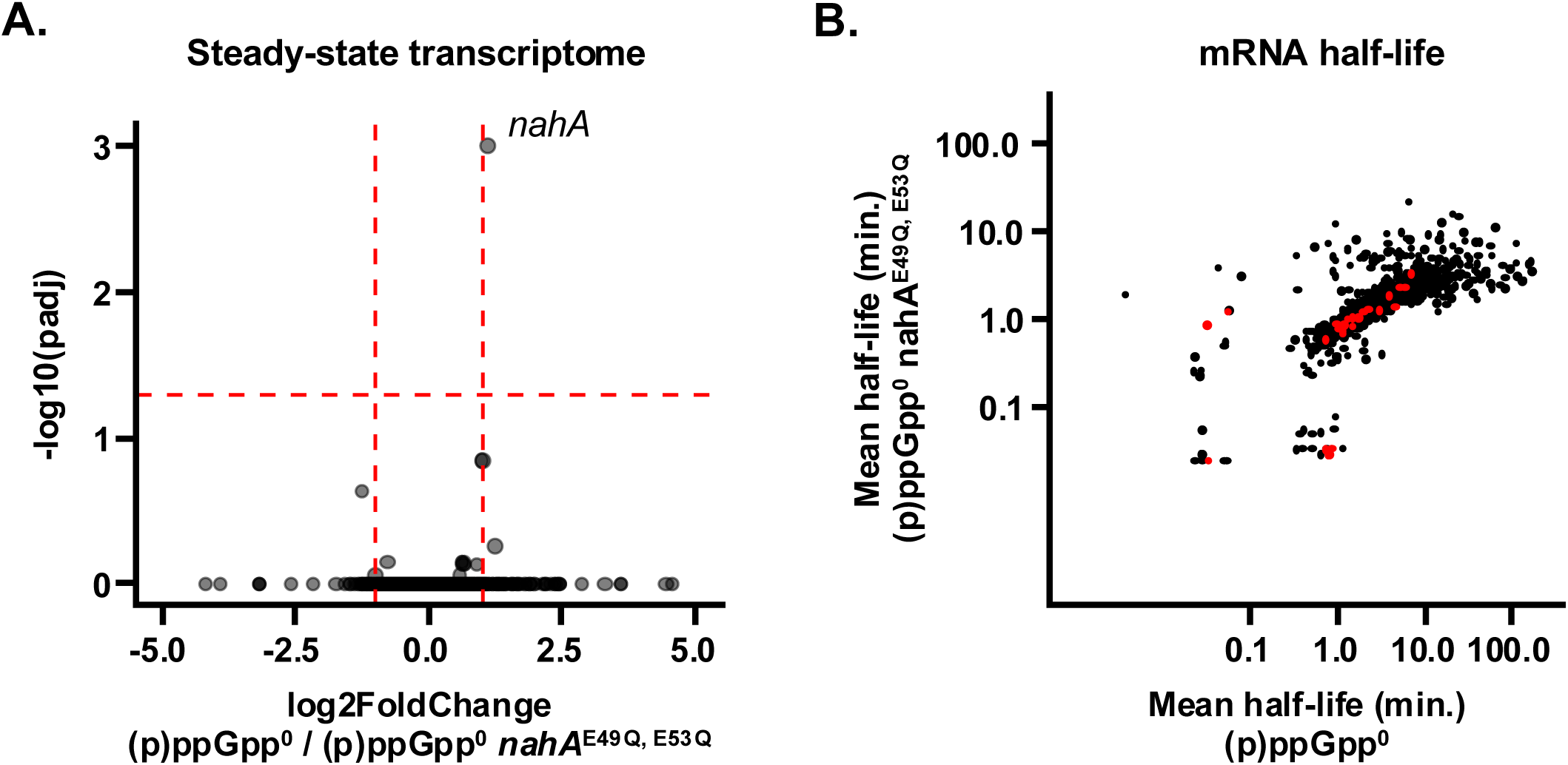
**(A)** NahA does not greatly affect the steady-state transcriptome. Each point represents one gene. Horizontal red-dashed line represents *p_adj_* = 0.05, and vertical red-dashed lines represent log_2_ fold changes = 1 and -1. (**B)** NahA does not greatly destabilize mRNA. Each point represents one gene, where the X-coordinate is the half-life in (p)ppGpp^0^ and the Y-coordinate is the half-life in (p)ppGpp^0^ *nahA^E49Q, E53Q^.* Red points are those that have *p* < 0.05 (Student’s T-test).

## Notes

### Competing Interest Statement

The authors have declared no competing interest.

