## Supplementary figures and images for "Starvation-independent alarmone production inhibits translation through GTP depletion"

### Supplemental dataset S2

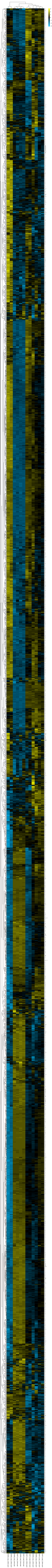
